# Clathrin-independent endocytosis and retrograde transport in cancer cells tune immune synapse organization and CD8 T cell response

**DOI:** 10.1101/2024.10.28.620627

**Authors:** Shiqiang Xu, Alix Buridant, Thibault Hirsch, Céline Duhamel, Benjamin Ledoux, Massiullah Shafaq-Zadah, Estelle Dransart, Louise Thines, Ludger Johannes, Pierre Van Der Bruggen, Pierre Morsomme, Henri-François Renard

## Abstract

Endophilin A3-mediated clathrin-independent endocytosis (EndoA3-mediated CIE) contributes to the internalization of immunoglobulin-like proteins, including key immune synapse components. Here, we identify ICAM1 as a novel EndoA3-dependent cargo, alongside ALCAM. We demonstrate that both proteins subsequently follow retromer-dependent retrograde transport to the *trans*-Golgi network (TGN) in cancer cells. From there, we propose that they undergo polarized redistribution to the plasma membrane, where they contribute to immune synapse formation between cancer cells and cytotoxic CD8 T cells. Disruption of EndoA3 or retromer components significantly affects the response of autologous cytotoxic CD8 T cells, as evidenced by reduced cytokine production and secretion, but increased lytic activity, while proliferation and later activation marker expression remain intact. This is accompanied by diminished ICAM1 density at the immune synapse, where we observe it arriving via polarized vesicular transport, indicating altered synapse organization. Indeed, cancer cells lacking EndoA3-mediated CIE or retromer form enlarged immune synapses that fail to sustain full T cell cytokine secretion, suggesting a compensatory attempt by T cells to overcome the defective synapse, while likely promoting more transient contacts that potentially favor serial killing. Together, these findings reveal that EndoA3-mediated CIE and retrograde transport act in concert in cancer cells to relocate immune synapse components via the Golgi, thereby fine-tuning the balance between cytotoxic T cell cytokine secretion and lytic activity. These insights contribute to a better understanding of the mechanisms governing immune synapse formation and organization, providing a necessary foundation for the long-term identification of new strategies to enhance T cell–mediated anti-tumor immunity.

**Significance Statement:** This study uncovers a novel mechanism by which clathrin-independent endocytosis (CIE) and retrograde transport collaborate to regulate immune synapse dynamics in cancer cells. We identify ICAM1 as a new cargo of Endophilin A3-mediated CIE, highlighting its role in the polarized redistribution of immune synapse components critical for cytotoxic CD8 T cell activation. By linking a specific CIE mechanism and retrograde transport to immune synapse function, our findings provide new insights into cancer-immunity interactions and suggest potential therapeutic strategies to enhance immune responses by targeting protein trafficking pathways.

## Introduction

Endocytosis is a fundamental physiological process involving membrane bound carrier-mediated internalization of extracellular substances and cell membrane components into the cytoplasm. This process is essential for maintaining plasma membrane homeostasis and regulating signal transduction. Broadly, endocytosis can be classified into conventional clathrin-mediated endocytosis (CME) and unconventional clathrin-independent endocytosis (CIE). While CME has been extensively characterized, CIE remains less understood and complex as it includes several distinct endocytic mechanisms (1, 2). Following endocytosis, internalized cargoes are sorted in early/sorting endosomes, where they are either directed toward lysosomes for degradation or recycled to the plasma membrane for further activity. Some are recycled directly to the plasma membrane via Rab4/Rab11 recycling endosomes (3, 4), while others are retrieved by retromer or Commander complexes for transport to the *trans*-Golgi network (TGN), a process termed retrograde transport (5–8). Increasing evidence has linked retrograde transport to the maintenance of cell polarity (9). For instance, β_1_ integrins are transported to the TGN via the retrograde transport route post-endocytosis, from where they are redistributed in a polarized manner to the leading edge of migrating cells, supporting front-rear polarity crucial for persistent migration (10). Similarly, in T cells, adaptor molecules such as the Linker for Activation of T cells (LAT) are transported to the TGN after endocytosis and redistributed to the immune synapse (11). The immune synapse is a polarized structure formed between immune and target cells (*e.g.*, cytotoxic CD8 T cells and cancer cells) (12), which exemplifies the intricate interplay between retrograde transport and cell polarity. Despite its importance, the connection between specific CIE mechanisms and retrograde transport in polarized cellular contexts remains largely underexplored.

Endophilin A (EndoA) proteins are key players in endocytosis, belonging to the BAR (Bin/Amphiphysin/Rvs) domain protein family known for their ability to sense and/or induce membrane curvature (13, 14). Mammalian cells express three isoforms of EndoA: EndoA1, A2, and A3 (15, 16). EndoA1 is predominantly expressed in the brain, EndoA3 is abundant in both the brain and testes, and EndoA2 is ubiquitously expressed across tissues (17, 18). Initially, EndoA proteins were implicated in CME in neuronal cells, where they recruit dynamins and synaptojanins via interactions between their Src-homology 3 (SH3) domains and proline-rich domains (PRDs), facilitating vesicle scission (19, 20) and subsequent clathrin uncoating (21, 22).

Recent studies have identified EndoA2 protein as a central player in clathrin-independent Fast Endophilin-Mediated Endocytosis (FEME) (23). EndoA2 has also been implicated in membrane scission events during the clathrin-independent uptake of Shiga and cholera toxins (24, 25). Intriguingly, the three EndoA isoforms are not fully functionally redundant in CIE. Only EndoA3, and not EndoA1 or A2, mediates the endocytosis of the immunoglobulin (Ig)-like protein ALCAM (Activated Leukocyte Cell Adhesion Molecule). The endocytosis of ALCAM, the first identified EndoA3-specific cargo, operates in an EndoA3-dependent manner and is driven by extracellular Galectin-8 and glycosphingolipids, in agreement with the glycolipid-lectin (GL-Lect) endocytosis mechanism (26–28). Another Ig-like protein, L1 Cell Adhesion Molecule (L1CAM, alternative name CD171), has since been confirmed as a cargo for EndoA3-mediated CIE (29).

ALCAM (CD166) is an adhesion molecule broadly expressed across tissues and overexpressed in various cancers, including bladder, prostate and colorectal carcinomas (30–32). ALCAM mediates cell-cell interactions either through homophilic binding with other ALCAM molecules (33) or by heterophilic binding with CD6 (34). CD6, a member of the scavenger receptor cysteine-rich (SRCR) protein family, is predominantly expressed in immune cells, particularly T lymphocytes, and in certain neuronal cells (35, 36). The ALCAM-CD6 interaction has been shown to play a critical role in immune synapse formation, providing essential co-stimulatory signals for optimal T cell activation and proliferation (37–39).

Given the polarized nature of the immune synapse, we hypothesized that EndoA3-mediated endocytosis in cancer cells could influence immune synapse formation through the subsequent retrograde transport and polarized redistribution of its components, such as ALCAM. Moreover, since both ALCAM and L1CAM are Ig-like proteins, we speculated that EndoA3-mediated endocytosis might preferentially facilitate the uptake of Ig-like proteins. Interestingly, Intercellular Adhesion Molecule 1 (ICAM1, alternative name CD54), a well-characterized immune synapse component critical for co-stimulating T cell activation via its interaction with Lymphocyte Function-associated Antigen 1 (LFA-1) on the T cells (40–42), is also an Ig-like protein. This prompted us to investigate whether ICAM1 serves as an EndoA3-dependent CIE cargo in cancer cells and whether its role in immune synapse formation relies on EndoA3-mediated CIE and retrograde transport.

Through a combination of cell biology and immunology experiments, we identify ICAM1 as a novel EndoA3-dependent endocytic cargo. We further show that both ALCAM and ICAM1 undergo retromer-dependent retrograde transport to the TGN following EndoA3-mediated internalization. Disruption of EndoA3-mediated endocytosis or retromer-dependent retrograde transport in cancer cells significantly affects the response of autologous cytotoxic CD8 T cells, leading to reduced cytokine production and secretion, while paradoxically enhancing their lytic activity. Because ICAM1 reaches the immune synapse through polarized vesicular transport, we propose that this phenotype arises from defective polarized redistribution of immune synapse components, including ICAM1, from the Golgi to the plasma membrane. Consistent with this hypothesis, we find that disrupting EndoA3-mediated endocytosis in cancer cells reduces ICAM1 recruitment to the immune synapse, likely reflecting compromised structural integrity and formation. Interestingly, depleting EndoA3 or VPS26A increases the size of immune synapses, which reduces ICAM1 density at the contact zone. This synapse enlargement may represent an attempt by T cells to compensate for the loss of correctly localized components. However, this adaptation is insufficient to fully sustain CD8 T cell cytokine production and secretion, suggesting impaired synapse maturation. Conversely, we observe enhanced lytic activity, suggesting that these enlarged immune synapses promote more rapid detachment and re-engagement of T cells, potentially facilitating “serial killing”. Collectively, our findings reveal an unexplored functional link between EndoA3-mediated clathrin-independent endocytosis and retrograde transport, which cooperate in cancer cells to relocate immune synapse components via the Golgi, thereby fine-tuning the balance between cytotoxic T cell cytokine secretion and lytic activity.

## Results

### ALCAM and ICAM1 are cargo clients of retromer-dependent retrograde trafficking

ALCAM and ICAM1 are known components of immune synapses (37–39, 41, 42). Given the polarized nature of these structures and the previously reported connection between retrograde transport and cell polarity (10, 11), we investigated whether ALCAM and ICAM1 serve as cargoes for retrograde transport. To address this question, we employed a SNAP-tag-based assay to study retrograde transport of cargoes from the plasma membrane to the Golgi apparatus (43) (Fig. 1A). Briefly, benzylguanine (BG)-labelled cargo-specific antibodies (IgG) were incubated with cells stably expressing the Golgi-resident Galactosyltransferase protein (GalT) fused to GFP and SNAP-tag (GalT-GFP-SNAP). If the cargo follows the retrograde route after endocytosis, it will be delivered to the TGN along with the BG-labelled antibody. Upon reaching the TGN, BG covalently reacts with the SNAP-tag, generating a large complex (IgG-SNAP-GFP-GalT), which can be immunopurified and analyzed by Western blot.

**Figure 1.**
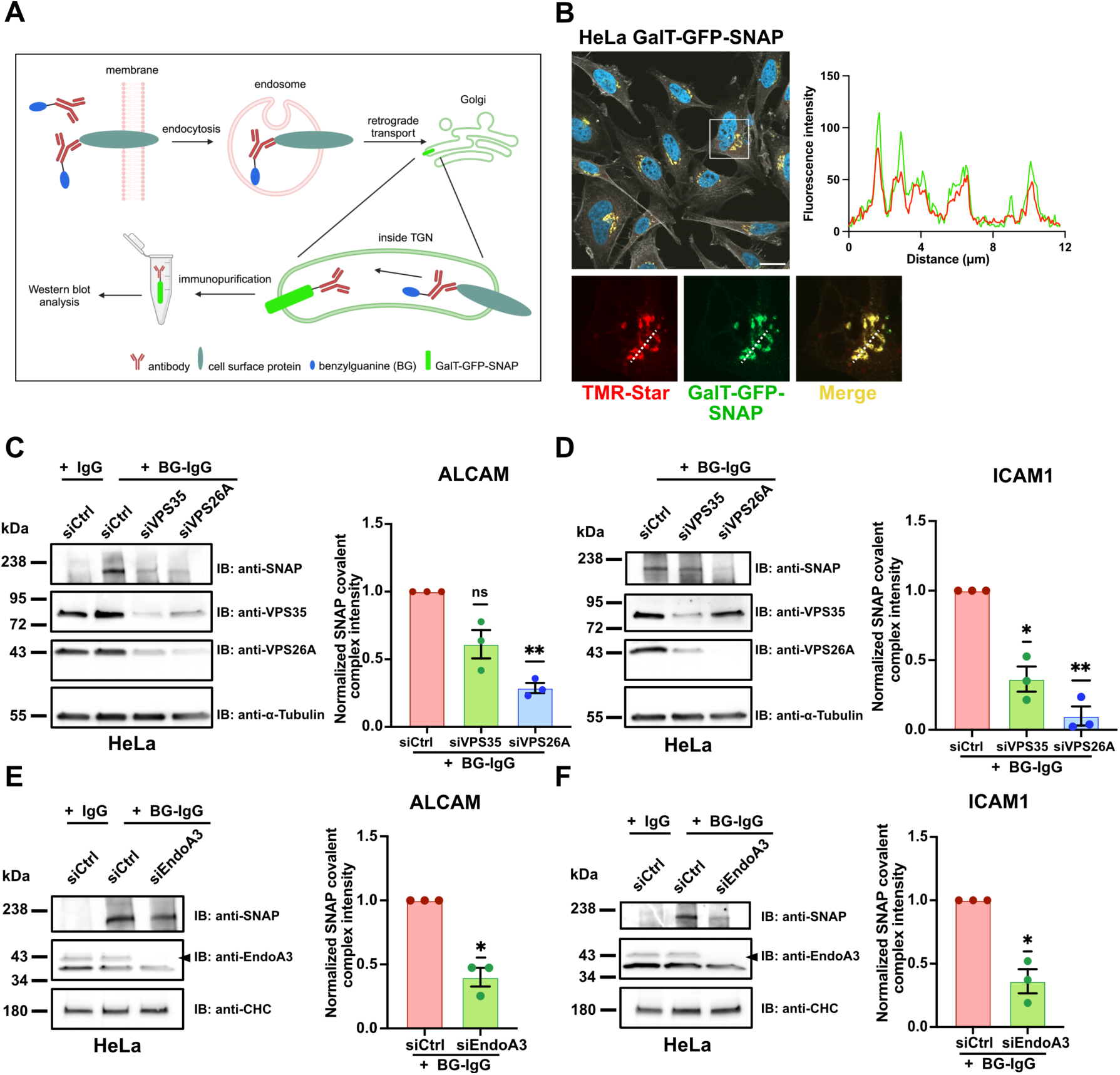
Immune synapse components ALCAM and ICAM1 are retrograde transport cargoes that rely on EndoA3 and retromer. A Illustration of the SNAP-tag-based BG-labelled antibody uptake assay to study membrane protein endocytosis and retrograde transport. B Confocal images of GalT-GFP-SNAP (green) and TMR-Star (red) in HeLa cells stably expressing the Golgi-resident GFP-fused SNAP-tag construct (HeLa GalT-GFP-SNAP). Actin (phalloidin, white) and nuclei (DAPI, blue) were also stained. Fluorescence intensity profile was made along the dashed line region in enlarged cropped area, and shows the colocalization of both signals. Scale bar: 20 μm. C-F Retrograde transport of ALCAM and ICAM1. Continuous BG-labelled anti-ALCAM (C and E) and anti-ICAM1 (D and F) antibody uptake for 4h at 37°C in HeLa GalT-GFP-SNAP cells. (C-D) Western blot analysis of HeLa GalT-GFP-SNAP cells transfected for 72h with siRNAs: negative control (siCtrl) or against retromer subunits (siVPS35 and siVPS26A). Immunodetection made with anti-SNAP, anti-VPS35, anti-VPS26A and anti-α-Tubulin (loading control) antibodies. Quantification of the covalent IgG-SNAP-GFP-GalT complex is shown as fractions of siCtrl condition (histogram). Quantification of VPS35 and VPS26A depletion is shown in Fig. S1C. (E-F) Western blot analysis of HeLa GalT-GFP-SNAP cells transfected for 72h with siRNAs: negative control (siCtrl) or against EndoA3 (siEndoA3). Immunodetection made with anti-SNAP, anti-EndoA3 and anti-clathrin heavy chain (CHC, loading control) antibodies. Quantification of the covalent IgG-SNAP-GFP-GalT complex (IB:anti-SNAP) is shown as fractions of siCtrl condition (histogram). Quantification of EndoA3 depletion is shown in Fig. S1H. C Data information: In (B), images are from a single experiment. Quantification data (C-F) are pooled from three independent experiments. Data are presented as mean ± SEM. *P < 0.05, **P < 0.01. One sample *t* and Wilcoxon test. Source data are available online for this figure.

We used a HeLa cell line stably expressing the Golgi-resident SNAP-tag construct (HeLa GalT-GFP-SNAP), validated in earlier studies (10, 44). Additionally, we generated a LB33-MEL cell line stably expressing the same Golgi-resident SNAP-tag construct (LB33-MEL GalT-GFP-SNAP, Fig. S1A-B). Of note, the LB33-MEL melanoma cell line was derived from a melanoma patient (LB33) in 1988. A specific autologous CD8 T cell line targeting this melanoma, obtained from the same patient, enables *in vitro* reconstitution of immune synapses (45). First, we validated the SNAP-tag functionality in both cell lines using the SNAP-Cell^®^ TMR-Star reagent (Fig. 1B for HeLa; Fig. S1A for LB33-MEL). We observed a strong and specific colocalization of TMR-Star with the GFP signal of the SNAP-tag constructs in both cell lines. Second, immunofluorescence experiments showed that the SNAP-tag construct co-distributes with the *TGN*-specific marker TGN46 in LB33-MEL cells, confirming its correct and specific localization (Fig. S1B).

The retromer complex, composed of VPS26, VPS29 and VPS35 subunits in mammalian cells, is the best-studied machinery for retrograde transport (5, 46). Two paralogues of VPS26 subunit exist: VPS26A, expressed in most tissues, and VPS26B, mainly enriched in the brain (47, 48). To determine whether ALCAM and ICAM1 are cargoes of the canonical retromer complex, we conducted SNAP-tag-based retrograde transport assays in HeLa and LB33-MEL cell lines where VPS35 or VPS26A was knocked down by RNA interference (Fig. 1C-D, S1C for HeLa; Fig. S1D-E for LB33-MEL). BG-labelled anti-ALCAM or anti-ICAM1 antibodies were added to the cell culture medium for a 4-hour continuous uptake and the production of a covalent IgG-SNAP-GFP-GalT complex was monitored by Western blot (simplified as SNAP signal).

We observed that both ALCAM and ICAM1 are retrograde transport cargoes in cells treated with negative control siRNA, as IgG-SNAP-GFP-GalT complexes were detected by Western blot after incubation with BG-labelled antibodies (band ∼ 210 kDa; Fig. 1C-D for HeLa; Fig. S1D for LB33-MEL). As a negative control, unlabeled antibodies (IgGs) showed no detectable SNAP signal in Western blot (Fig. 1C). Interestingly, depletion of VPS35 or VPS26A caused a significant reduction in the SNAP signal for both cargoes (Fig. 1C-D, S1C for HeLa; Fig. S1D-E for LB33-MEL). Of note, VPS26A depletion resulted in a stronger decrease in retrograde transport efficiency compared to VPS35 (Fig. 1C-D).

To explore additional factors involved in retrograde transport, we examined the role of Rab6 GTPase, a critical molecular player in secretion which controls the formation of membrane tubules originating from the TGN (49, 50). Disrupting Rab6 indirectly inhibits retrograde transport to the TGN of cargoes requiring subsequent polarized anterograde redistribution (10). Interestingly, Rab6 depletion impaired retrograde transport of ALCAM to the TGN, showing effects similar to retromer component depletion (Fig. S1F-G).

Together, these data demonstrate that retromer-mediated retrograde transport is critical for trafficking ALCAM and ICAM1 to the Golgi and that this process requires efficient secretion from the TGN (as evidenced by the involvement of Rab6).

Importantly, previous studies from our lab demonstrated that the endocytosis of ALCAM is mediated by EndoA3-dependent CIE (26, 51). We therefore investigated whether EndoA3 depletion could affect the post-endocytic retrograde transport of ALCAM to the TGN. To test this, we depleted EndoA3 in HeLa cells using RNA interference (Fig. 1E-F, S1H) and observed 60% of reduction in the IgG-SNAP signal intensity (Fig. 1E). Additionally, we performed retrograde transport assays with BG-labelled anti-ALCAM antibodies in the SNAP-tag-expressing LB33-MEL cell line (Fig. S1I). Since LB33-MEL cells do not express detectable levels of EndoA3 as confirmed by Western blot (26), we transiently transfected these cells with an EndoA3-GFP expression plasmid. Remarkably, EndoA3 expression increased the SNAP signal by ∼50% compared to the control condition where cells were transfected with a plasmid expressing free GFP (Fig. S1I). Given that ICAM1 is also a member of the Ig-like protein family, we investigated whether the retrograde transport of this cargo depends on EndoA3. Interestingly, EndoA3 depletion in HeLa cells resulted in a significant reduction of the ICAM1 IgG-SNAP signal by 64% (Fig. 1F), comparable to the effect observed for ALCAM (Fig. 1E).

Collectively, these results suggest that: (i) ICAM1 is potentially a novel EndoA3-dependent cargo, (ii) substantial fractions of ALCAM and ICAM1 are transported to the TGN in a retromer-dependent manner after EndoA3-mediated CIE, and (iii) introducing EndoA3 in EndoA3-negative cells enhances the retrograde transport of these cell adhesion molecules to the TGN. These findings underscore a tight functional link between a specific clathrin-independent endocytic mechanism – mediated by EndoA3 – and the subsequent retrograde transport of cargoes to the TGN.

### EndoA3 expression enhances the internalization of immune synapse components ALCAM and ICAM1 in cancer cells

To further confirm that ICAM1 is an EndoA3-dependent cargo, we employed live-cell TIRF imaging to directly observe the dynamics of EndoA3 and ICAM1 at the plasma membrane of both HeLa and LB33-MEL cell lines stably expressing EndoA3-GFP. Of note, the EndoA3-GFP-expressing HeLa cell line was generated and validated in earlier studies from our lab (26, 29), while the EndoA3-GFP-expressing LB33-MEL cell line (hereafter referred to as LB33-MEL EndoA3+) was developed specifically for this study. Both cell lines were then transiently transfected to express ICAM1-mScarlet.

Interestingly, EndoA3-GFP formed circular structures around ICAM1 patches in both cell lines (Fig. 2A-B, S2A-B, white arrows; Movie S1-2, S5-6), and ICAM1 signals gradually disappeared from the plasma membrane (Fig. 2A-B, S2A-B; Movie S1-2, S5-6). In addition, ICAM1-positive punctate structures were observed, where EndoA3-GFP was recruited before both protein signals disappeared from the cell surface (Fig. 2A-B, S2B, kymographs; Movie S3-4 and S7), indicating active endocytic events. This behavior mirrors TIRF observations previously reported for other EndoA3-dependent endocytic cargoes, such as ALCAM and L1CAM (26, 29). Collectively, these observations confirm that ICAM1 is a novel EndoA3-dependent endocytic cargo.

**Figure 2.**
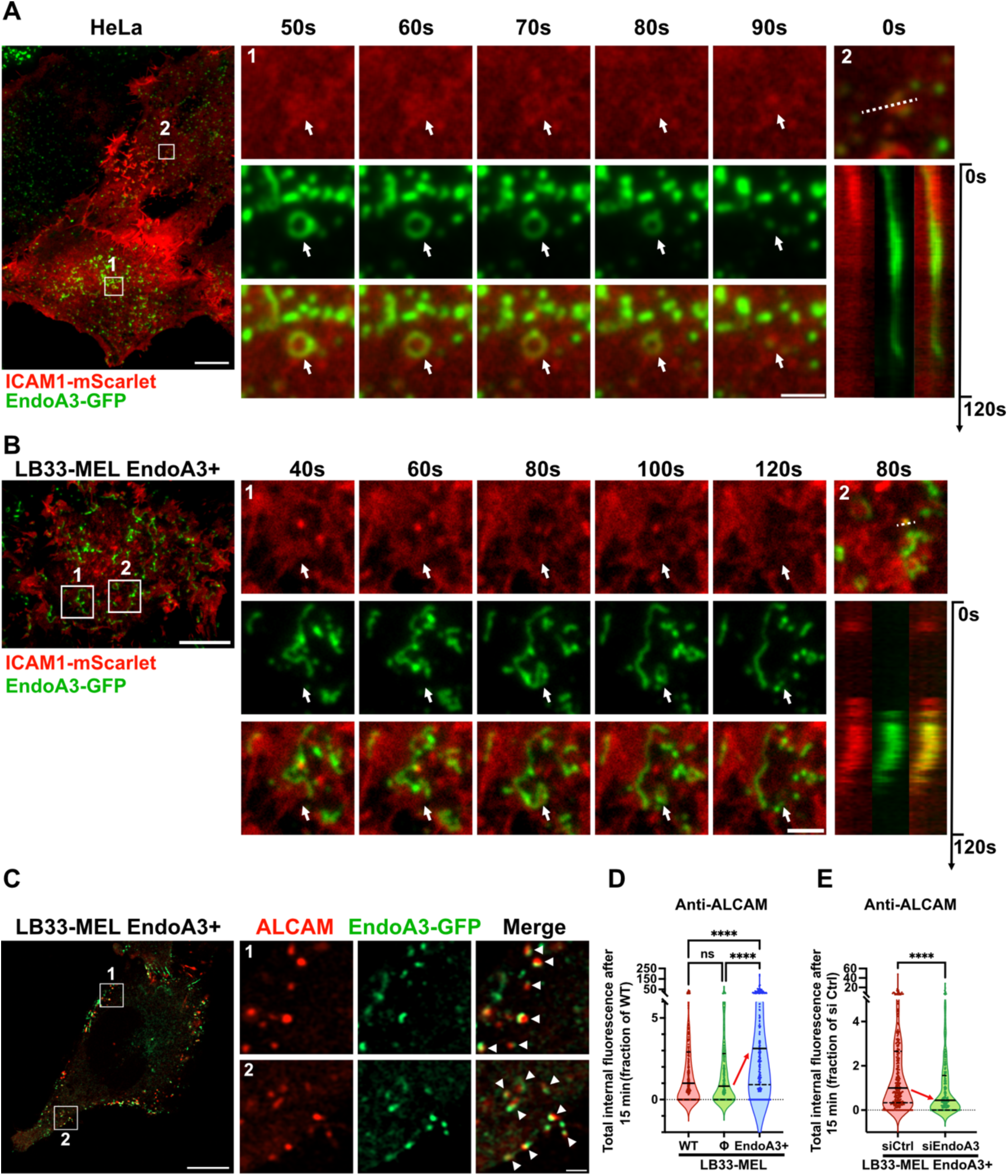
EndoA3-dependent CIE mediates the uptake of immune synapse components ALCAM and ICAM1 in cancer cells. A-B Live-cell TIRF images of EndoA3-GFP (stable) and ICAM1-mScarlet (transient) in HeLa (A) and LB33-MEL (B) cells. Time series show enlarged cropped areas corresponding to region 1 in the full-size images, and are extracted from Movie S1 (A) and Movie S2 (B). White arrows indicate dynamic co-distribution of both signals. Kymographs were made along the dashed lines in the enlarged cropped areas corresponding to region 2 (A-B; Movie S3-S4). Scale bars: 10 μm (full size image), 2 μm (enlarged cropped areas). C-E Continuous uptake of anti-ALCAM antibody for 15 min at 37°C in the following LB33-MEL cell lines: wild-type (WT, D), stably transfected with empty plasmid (Φ, D), or stably expressing EndoA3-GFP (LB33-MEL EndoA3+, C-E). In (E), cells were transfected with siRNAs: negative control (siCtrl) or against EndoA3 (siEndoA3). Quantification of EndoA3 depletion by western blots in Fig. S3C. (C) Airyscan images of Anti-ALCAM (red) and EndoA3-GFP (green). White arrowheads show colocalization between ALCAM and EndoA3. Scale bars: 10 μm (full size image), 1 μm (enlarged cropped areas). (D-E) Quantifications of anti-ALCAM internalization, expressed as fractions of WT condition (D) or siCtrl condition (E). (D) n cells: WT, n = 270; Φ, n = 279; EndoA3+, n = 274. (E) n cells: siCtrl, n = 350; siEndoA3, n = 234. Representative image examples in Fig. S3A-B. Data information: All images (A-C) are representative of two independent experiments. In (D-E), data are pooled from three independent experiments. Data are presented as median and quartiles. ns, not significant. ****P < 0.0001 (D, Kruskal-Wallis test with Dunn’s multiple comparison test; E, Mann-Whitney test). Source data are available online for this figure.

To investigate the role of EndoA3-dependent CIE and subsequent retrograde transport in cancer cells on the response of cytotoxic CD8 T lymphocytes, we further used the LB33-MEL cellular model. This cell line expresses a mutated antigenic peptide encoded by the *MUM3* gene. The MUM-3 peptide is presented on the cell surface by HLA-A*68012 molecules, an HLA-A28 subtype specific to patient LB33 (45, 52). A CD8 T cell line targeting the MUM-3 peptide, derived from the same patient in 1990 (45), can form immune synapses with LB33-MEL cells *in vitro*. Although LB33-MEL cells lack detectable levels of EndoA3 (26), we have observed above that exogenous expression of EndoA3 enhances the retrograde transport of cell adhesion molecules (Fig. S1I).

Consistent with this result, high resolution Airyscan confocal images showed a clear colocalization between EndoA3-GFP and ALCAM, after a 15-minute uptake of anti-ALCAM antibodies in LB33-MEL EndoA3+ cells (Fig. 2C, white arrowheads). This observation confirmed further that this cell line recapitulates EndoA3-mediated endocytosis. Additional quantitative analyses revealed that the uptake of anti-ALCAM antibodies was three times higher in LB33-MEL EndoA3+ cells compared to wild-type or empty plasmid-transfected LB33-MEL cells (Fig. 2D, S3A). In addition, we performed a reverse-rescue uptake assay in LB33-MEL EndoA3+ cells transfected with EndoA3 siRNAs (Fig. 2E, S3B). ALCAM uptake was significantly reduced by 55% upon EndoA3 knockdown (Fig. 2E). Of note, exogenously expressed EndoA3-GFP protein was efficiently knocked down by siRNAs in LB33-MEL EndoA3+ cells, with an 80% reduction as shown by immunoblotting (Fig. S3C).

Taken together, these findings confirm that LB33-MEL EndoA3+ cells recapitulate functional EndoA3-dependent CIE, enhancing the uptake of canonical EndoA3-dependent cargoes. This cell line provides a robust model to assess the effects of EndoA3-mediated CIE and subsequent retrograde transport on CD8 T cell response.

### EndoA3-dependent CIE and retromer-mediated retrograde transport in cancer cells modulate cytotoxic CD8 T cell cytokine secretion and lytic activity

Based on our results and considering that ALCAM and ICAM1 are key components of immune synapses, we hypothesized that their clathrin-independent endocytosis and subsequent retrograde transport might influence CD8 T cell response. To test this hypothesis, we conducted CD8 T cell stimulation assays using LB33-MEL cells, in the presence or absence of EndoA3. Specifically, LB33-MEL EndoA3+ cells were treated with either control or EndoA3-targeting siRNAs before co-culture with CD8 T cells. Intracellular cytokine production in T cells was then assessed by flow cytometry (Fig. 3A).

**Figure 3.**
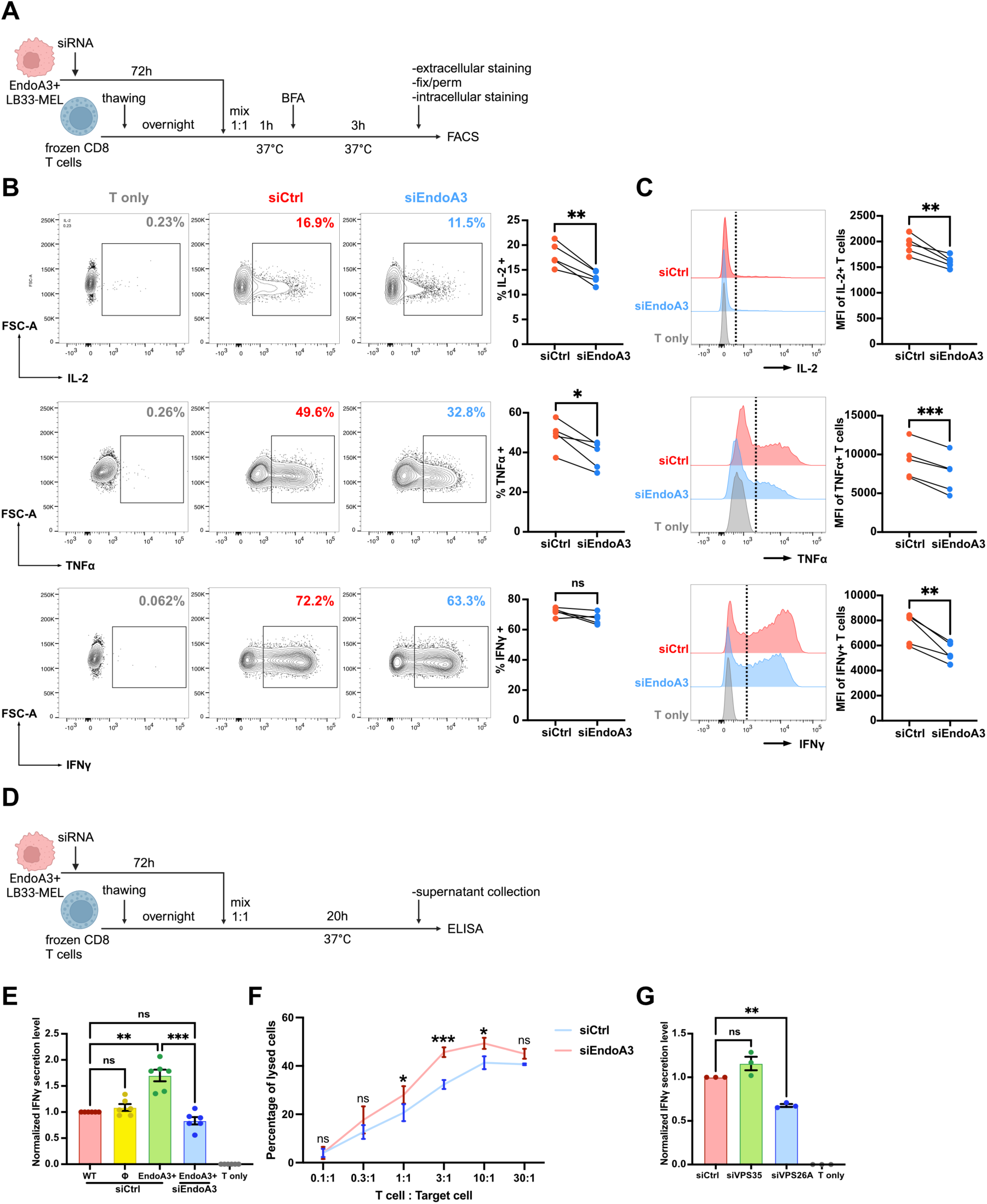
Inhibition of EndoA3-dependent endocytosis and retrograde transport impairs T cell activation but increases their lytic activity. A Scheme of flow cytometry analysis of cytokine production inside CD8 T cells stimulated by siRNA transfected LB33-MEL EndoA3+ cells. BFA, Brefeldin A. B-C Flow cytometry analysis of CD8 T cell intracellular cytokine production after co-culture for 4h at 37°C with: no LB33-MEL (T only), LB33-MEL EndoA3+ transfected with negative control siRNA (siCtrl) or with siRNA against EndoA3 (siEndoA3). Quantification of EndoA3 depletion is shown in Fig. S4A. (B) Representative examples and quantifications (scatter plots) of the percentages of cytokine producing CD8 T cells (up, IL-2; middle, TNFα; bottom, IFNγ) after being stimulated by siCtrl or siEndoA3 treated LB33-MEL EndoA3+ cells. (C) Representative examples and quantifications (scatter plots) of the absolute amount of cytokines produced by CD8 T cells (up, IL-2; middle, TNFα; bottom, IFNγ; presented as median fluorescence intensity, MFI) after stimulation by siCtrl or siEndoA3 treated LB33-MEL EndoA3+ cells. D Scheme of ELISA analysis of cytokine secretion from CD8 T cells stimulated with siRNA transfected LB33-MEL EndoA3+ cells. E Quantification of IFNγ secretion (detected by ELISA) from CD8 T cells, cultured alone (T only) or co-cultured for 20h with the following LB33-MEL cell lines: wild-type LB33-MEL cells (WT), LB33-MEL cells stably transfected with empty (Φ) or EndoA3-GFP encoding plasmid (EndoA3+), treated with negative control siRNA (siCtrl) or with EndoA3-targeting siRNA (siEndoA3). Data are presented as fractions of WT siCtrl condition. The absolute concentration of secreted IFNγ in the supernatant of WT siCtrl condition is 2203 ± 201 (mean ± SEM) pg/mL. F Quantification of CD8 T cell cytolytic efficiency against LB33-MEL EndoA3+ cells transfected with negative control siRNA (siCtrl) or siRNA against EndoA3 (siEndoA3). CD8 T cell killing efficiency was determined by Chrome 51 release assay and is presented as percentage of lysed LB33-MEL EndoA3+ cells. Different T cell:Target cell ratios were tested. G Quantification of IFNγ secretion (detected by ELISA) from CD8 T cells, cultured alone (T only) or co-cultured for 20h at 37°C with EndoA3+ LB33-MEL cells transfected with siRNAs: negative control (siCtrl), or against retromer subunits (siVPS35 or siVPS26A). Data are presented as fractions of siCtrl condition. The absolute concentration of secreted IFNγ in the supernatant of siCtrl condition is 4077 ± 99.62 (mean ± SEM) pg/mL. Quantification of VPS35 and VPS26A depletion is shown in Fig. S4L. Data information: In (B-C), data are pooled from five independent experiments. In (E), data are pooled from six independent experiments. In (F-G), data are pooled from three independent experiments. In (E,G), data are presented as mean ± SEM. ns, not significant. *P < 0.05, **P < 0.01, ***P < 0.001 (B and C, paired *t* test; E, RM one-way ANOVA with Tukey’s multiple comparison test; F, two-way ANOVA with Sidak’s multiple comparison test; G, RM one-way ANOVA with Dunnett’s multiple comparison test). Source data are available online for this figure.

When CD8 T cells were stimulated with EndoA3-depleted LB33-MEL cells, we observed a significant decrease in both the percentage of T cells producing interleukin-2 (IL-2) and tumor necrosis factor α (TNFα) (Fig. 3B, upper and middle panels), as well as the absolute amounts of these cytokines (Fig. 3C, upper and middle panels), compared to CD8 T cells stimulated with LB33-MEL EndoA3+ cells. Although the percentage of T cells producing interferon γ (IFNγ) remained unchanged (Fig. 3B, bottom panel), the absolute amount of IFNγ produced was still significantly reduced (Fig. 3C, bottom panel). Of note, the efficiency of EndoA3-GFP depletion was confirmed by flow cytometry (Fig. S4A).

Next, we assessed IFNγ secretion by CD8 T lymphocytes in co-culture supernatants using enzyme-linked immunosorbent assay (ELISA) (Fig. 3D). Wild-type, empty plasmid control, and EndoA3-GFP-expressing LB33-MEL cells all stimulated IFNγ secretion by CD8 T cells, compared to the control condition where T cells were cultured alone (Fig. 3E). Notably, EndoA3 expression significantly enhanced CD8 T cell response, leading to a ∼70% higher IFNγ secretion compared to wild-type or empty plasmid–transfected cancer cells (Fig. 3E). This stimulatory effect was lost upon EndoA3 depletion by RNA interference (Fig. 3E). To exclude the possibility that these effects are due to alterations in the biophysical properties of cancer cells following siRNA treatment (*i.e.* membrane tension), we analyzed their spreading area (Fig. S4B), aspect ratio (Fig. S4C), and roundness (Fig. S4D). None, or only minimal, changes were detected, indirectly indicating that the observed differences in T cell activation were likely not attributable to variations in cancer cell stiffness. In addition, to exclude the possibility that these effects are due to drastic changes of ICAM1 and ALCAM abundance at the plasma membrane, we measured their surface level by flow cytometry in cancer cells expressing or depleted for EndoA3 (Fig. S4E-F). We observed only minor variations which cannot account for the very significant drop in IFNγ secretion upon EndoA3 depletion.

To further dissect the mechanisms underlying EndoA3-mediated modulation of T cell responses, we examined additional parameters of CD8 T cell activation. We first analyzed surface expression of activation markers, including PD-1, CD137 and Tim-3, on CD8 T cells co-cultured with EndoA3-expressing or -depleted LB33-MEL cells (Fig. S4G-I). None of these markers were significantly altered by EndoA3 knockdown. Similarly, EndoA3 depletion did not affect T cell proliferation (Fig. S4J) nor degranulation (Fig. S4K). Unexpectedly, chromium release assays revealed that CD8 T cells exhibited increased cytotoxic activity when co-cultured with EndoA3-depleted cancer cells, approximately 15% higher at a 3:1 T cell–to–target cell ratio, corresponding to the maximal effect observed (Fig. 3F). Although seemingly paradoxical, this finding may align with a model in which EndoA3 depletion leads to disturbed immune synapse organization, thereby promoting shorter contacts with CD8 T cells and enhancing serial killing capacity. Shorter contacts with cancer cells may also explain the concomitant decrease in IFNγ secretion under the same conditions (see Discussion).

Together with our previous results, these observations support the hypothesis that EndoA3-mediated CIE in cancer cells influences retrograde transport and the subsequent polarized redistribution of immune synapse components, such as ALCAM and ICAM1, thereby affecting T cell response. To further test this model, we disrupted the retromer complex in LB33-MEL EndoA3+ cells by depleting the subunits VPS26A and VPS35 using RNA interference, and assessed the effects on CD8 T cell activation by measuring cytokine secretion. While VPS35 depletion had no significant impact, loss of VPS26A markedly impaired IFNγ secretion (Fig. 3G, S4L).

In summary, our data demonstrate that EndoA3-dependent endocytosis in cancer cells contributes to CD8 T cell response by promoting retrograde transport of immune synapse components such as ALCAM and ICAM1. This process likely enables their polarized redistribution from the TGN to the cell surface, facilitating efficient immune synapse assembly. Conversely, EndoA3 depletion in target cells reduces CD8 T cell cytokine production and secretion, but increases their lytic activity, reflecting changes in immune synapse organization and potentially enhanced serial killing dynamics driven by more transient immune synapse interactions.

### Inhibition of EndoA3-dependent endocytosis and retrograde transport affects ICAM1 recruitment in the vicinity of the contact zone with CD8 T cells

Building on the data presented above, effective EndoA3-dependent endocytosis tunes CD8 T cell response, possibly through an increased retrograde flux to the TGN and the subsequent polarized redistribution of immune synapse components to the plasma membrane. According to this hypothesis, depletion of EndoA3 or retromer subunits should perturb the recruitment of components to the immune synapse. Consequently, the structural features of the immune synapses would likely be affected. To test this, we monitored the recruitment of ICAM1 to the immune synapse using spinning-disk and high resolution Airyscan confocal microscopy, given the well-established role of ICAM1 in T cell activation (40).

While we were able to transiently transfect LB33-MEL cells with an ICAM1-expressing plasmid (Fig. 2B), the transfection efficiency was low, thereby limiting the possibility of using LB33-MEL cells for immune synapse imaging. To overcome this, we generated a stable HLA-A*68012-expressing HeLa cell line. Wild-type HeLa cells lack HLA-A*68012 and are therefore unable to present the MUM-3 antigenic peptide to stimulate CD8 T cell activation (Fig. S5A). Upon addition of exogenous MUM-3 peptide, the HLA-A*68012-expressing HeLa cells successfully stimulated anti-MUM-3 CD8 T cells to secrete IFNγ (Fig. S5A) and proliferate (Fig. S5B). Thereby, these data confirmed the functionality of this HLA-A*68012-expressing HeLa cell line.

To monitor the recruitment of ICAM1 to the immune synapse, HLA-A*68012-expressing HeLa cells transiently expressing EGFP-tagged ICAM1 were loaded with MUM-3 peptide and co-cultured with CD8 T cells. Using high-speed spinning-disk confocal microscopy, we observed a flux of ICAM1-positive tubulo-vesicular carriers emerging from the perinuclear region (corresponding to the Golgi area) and moving toward the contact site with the CD8 T cell (Fig. 4A; Movie S8), with fusion events of carriers occurring at the developing immune synapse-like conjugate (Fig. 4B; Movie S9). AI-based segmentation and tracking analyses showed that ICAM1-positive carrier trajectories were predominantly oriented toward the forming immune synapse (Fig. 4C), whereas carriers moving toward other cellular regions were markedly less frequent (Fig. 4D-E), as revealed by the quantification of track densities (Fig. 4F) and track proportions in the different directions (Fig. 4G) in these different regions. Aligning with these observations, immunofluorescence experiments using HeLa cells transiently expressing ICAM1-mScarlet revealed a strong accumulation of ICAM1 signal in the contact zone with CD8 T cells (Fig. S5C, white arrowheads). Altogether, these results provide direct evidence for polarized ICAM1 transport via vesicular trafficking toward the immune synapse.

**Figure 4.**
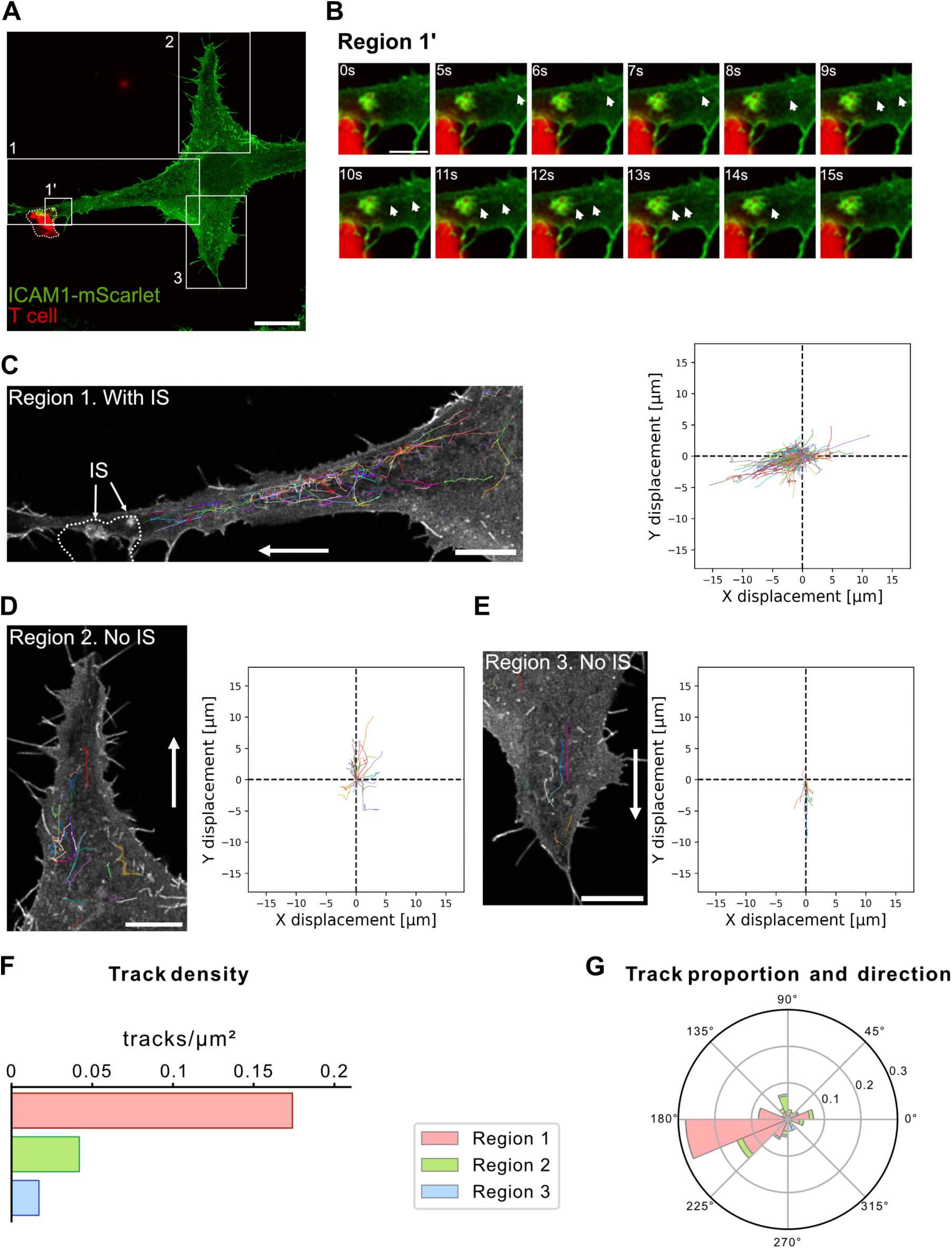
ICAM1 is directionally transported to the immune synapse via vesicular trafficking in cancer cells. A Representative live-cell spinning-disk confocal image of an immune synapse-like conjugate formed between a CD8 T cell (red) and an adherent, stable HLA-A*68012–expressing HeLa cell transiently expressing ICAM1-mScarlet (green) (Movie S8). Three regions were selected for quantitative analysis: region 1 at the T cell contact site, and regions 2 and 3 away from the conjugate (quantified in panels C-G). The T cell is delineated by a dotted line. Scale bar, 20 μm. B Time series of the enlarged cropped area (region 1′) from panel A, extracted from Movie S8 and shown in Movie S9. White arrowheads indicate ICAM1-positive tubulo-vesicular carriers moving toward the contact area and fusing with the conjugate membrane. Scale bar, 5 μm. C-E Tracking of ICAM1-positive carriers in the three regions defined in panel A-region 1 (C, with immune synapse) and regions 2-3 (D-E, without immune synapse). For each region, all recorded tracks are displayed on the cell image (left) and in corresponding X–Y displacement plots (right). Scale bar, 10 μm. F Quantification of track density in each region. Track density is markedly higher in region 1 where is the CD8 T cell contact. G Quantification of track directionality and proportion across regions. A higher proportion of tracks is oriented toward the T cell contact site in region 1. Data information: Images and quantifications are from a single movie, representative of two independent experiments.

Next, we treated ICAM1-mScarlet-expressing HeLa cells with control or EndoA3 siRNAs and incubated them with CD8 T cells (Fig. 5A-B, S5D-E). To facilitate quantification, this experiment was conducted on cells in suspension, which adopt a regular spherical shape, making them easier to segment using quantification algorithms (Fig. 5A, S5D, phalloidin channel). We measured the ICAM1 recruitment ratio in the vicinity of T cells (see Materials and Methods). Surprisingly, the ICAM1 signal was not confined to the contact region of the conjugates, but often expanded around the entire T cell (Fig. 5A, S5D). Quantifications revealed that depletion of EndoA3 in HLA-A*68012-expressing HeLa cells caused a slight but significant decrease in ICAM1 recruitment to the vicinity of anti-MUM-3 CD8 T cells (Fig. 5B), consistent with the previously observed impaired T cell cytokine secretion (Fig. 3).

**Figure 5.**
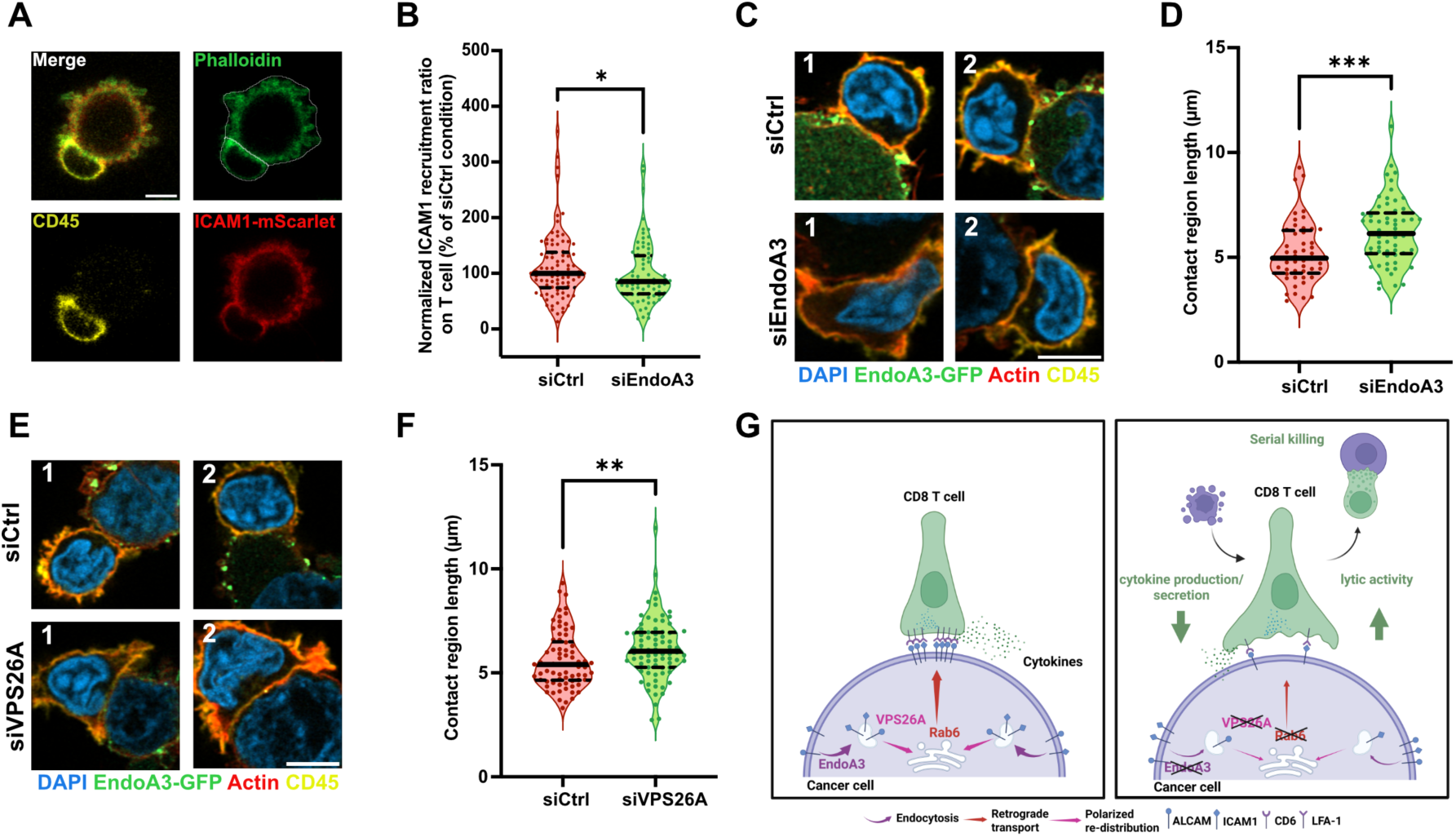
Inhibition of EndoA3-mediated endocytosis and retrograde transport affects ICAM1 recruitment to and structure of immune synapses. A Airyscan images of an immune synapse-like conjugate formed between a CD8 T cell (stained for CD45, yellow) and a stable HLA-A*68012-expressing HeLa cell (stained for actin, green) transiently expressing ICAM1-mScarlet (red) in suspension. Cell segmentation (white contour) was based on the actin staining for further quantifications (B). Scale bar: 5 μm. B Quantification of relative ICAM1 recruitment to the vicinity of CD8 T cell when an immune synapse-like conjugate is formed between a CD8 T cell and a stable HLA-A*68012-expressing HeLa cell transiently expressing ICAM1-mScarlet in suspension. HLA-A*68012-expressing HeLa cells were transfected for 72h with siRNAs: negative control (siCtrl) or against EndoA3 (siEndoA3). Western blot analysis of EndoA3 depletion is shown in Fig. S5E. n conjuagates: siCtrl, n = 85; siEndoA3, n = 75. C,E Airyscan images of immune synapse-like conjugates formed between CD8 T cells (stained for CD45, yellow) and stable EndoA3-GFP-expressing LB33-MEL cells (EndoA3-GFP, green) transfected for 72h with different siRNAs: negative control (siCtrl), siRNAs against EndoA3 (siEndoA3, C) or against VPS26A (siVPS26A, E). Actin (phalloidin, red) and nuclei (DAPI, blue) were also stained. Two examples are displayed per condition. Scale bar: 5 μm. D,F Quantifications of the sizes of immune synapse-like conjugates formed between CD8 T cells and stable EndoA3-GFP-expressing LB33-MEL cells transfected with different siRNAs from images in panels C and E. Western blot analyses of EndoA3 and VPS26A depletion are shown in Fig. S6C and S6D. (D) n conjugates: siCtrl, n = 48; siEndoA3, n = 58. (F) n conjugates: siCtrl, n = 74; siVPS26A, n = 78. G Working model. Immune synapse components ALCAM and ICAM1 are endocytosed into cancer cells through EndoA3-mediated CIE and are subsequently transported to the *TGN* in a retromer-dependent retrograde manner. Upon immune synapse formation, immune synapse components, such as ALCAM and ICAM1, are efficiently recruited to the contact area with CD8 T cells, likely from the *TGN* by polarized re-distribution, which can stabilize the immune synapse and promote cytokine production/secretion by CD8 T cells (left). In contrast, disruption of EndoA3-mediated CIE or retromer-dependent retrograde transport in cancer cells impairs the efficient recruitment of immune synapse components to the contact area with CD8 T cells, resulting in a larger but less stable immune synapses. Although CD8 T cells attempt to compensate by expanding their contact areas with cancer cells, this is insufficient to support efficient cytokine production and secretion. However, the reduced stability of the immune synapse may facilitate more rapid detachment and re-engagement of CD8 T cells, thereby enhancing their lytic activity, potentially through increased serial killing (right). Data information: In (A), (C) and (E), images are representative of three independent experiments. In (B), (D) and (F), data are pooled from three independent experiments. Data are presented as median and quartiles. *P < 0.05, **P < 0.01, ***P < 0.001 (B, Kolmogorov-Smirnov test; D and F, Mann-Whitney test). Source data are available online for this figure.

We then explored the consequences of EndoA3 or VPS26A depletion in cancer cells on the morphology of immune synapses. To study this, we returned to the LB33-MEL EndoA3+ cell line and imaged immune synapses formed with the CD8 T cells using Airyscan confocal microscopy (Fig. 5C and 5E, S6A-B). This experiment was also performed on cells in suspension for ease of quantitative analysis. We observed that immune synapses formed between CD8 T cells and EndoA3-depleted or VPS26A-depleted LB33-MEL EndoA3+ cells were larger than those formed between CD8 T cells and control LB33-MEL EndoA3+ cells. Quantification of immune synapse sizes confirmed this observation, showing that the immune synapses were 24% and 12% larger following EndoA3 and VPS26A depletion, respectively (Fig. 5D and 5F). Of note, the efficiency of siRNA transfections was validated by Western blot (Fig. S6C-D).

Together, these results show that inhibiting CIE or retrograde transport in cancer cells reduces ICAM1 recruitment in the vicinity of the contact zones with CD8 T cells and alters the organization of immune synapses. The resulting synapses are enlarged and display lower ICAM1 density, confirming structural alterations that were indirectly suspected from earlier CD8 T cell activation experiments (Fig. 3 and S4). Indeed, despite their increased size, these synapses fail to promote efficient cytokine secretion – which usually require fully mature synapses – although higher lytic activity is observed (Fig. 3 and S4).

In conclusion, disrupting endoA3-mediated CIE and downstream retrograde transport in cancer cells impairs the recruitment of immune synapse components, such as ICAM1, likely by altering their polarized redistribution from TGN to the plasma membrane. As a result, immune synapses may be less mature, with potential consequences on CD8 T cell responses. CD8 T cells may attempt to compensate for reduced recruitment of synaptic components by expanding their contact area with cancer cells, though this remains insufficient to support efficient cytokine production and secretion. This condition might also promote more transient contacts of CD8 T cells (*i.e.* faster detachment and re-engagement), which may favor serial killing (Fig. 5G).

## Discussion

In this study, we discovered that ICAM1 is a new cargo of EndoA3-mediated CIE, expanding the list of known EndoA3-dependent cargoes, which previously included the Ig-like cell proteins ALCAM and L1CAM (26, 29). Interestingly, the first study by Boucrot *et al.* (2015) on Fast Endophilin-Mediated Endocytosis (FEME) reported that the internalization of ICAM1 was not affected when all the three EndoA isoforms were depleted. This discrepancy could be attributed to differences in the cell lines used. Boucrot *et al.* (2015) used a human T cell line (Kit255), in which EndoA3 might be poorly expressed (17, 53), while we used human cancer cell lines – HeLa cells with endogenous EndoA3 expression and the LB33-MEL melanoma cell line engineered to stably express EndoA3. Moreover, it is now widely accepted that FEME is primarily mediated by the EndoA2 isoform (2).

Notably, all three confirmed EndoA3-dependent endocytic cargoes – ALCAM, L1CAM and ICAM1 – are Ig-like cell adhesion molecules. This suggests the possibility of a common recognition pattern for EndoA3-mediated CIE. Unlike CME, where cargo recognition is typically mediated by adaptor protein binding to specific sorting signals in the cytosolic tails of the cargoes (54), the mechanisms for cargo recognition in CIE are just beginning to emerge (28, 55). In FEME, EndoA2 is thought to directly recognize and sort G protein-coupled receptors (GPCRs) by binding to PRDs in GPCRs through their SH3 domains (23, 56), or indirectly sort cargoes by interacting with other proteins, such as binding to CIN85 for EGFR and HGFR (57, 58), or srGAP1 for ROBO1 (59). However, how EndoA3-dependent endocytic cargoes are recognized and sorted remains unclear. Our lab has previously shown that the cytosolic tails of ALCAM and L1CAM physically interact with EndoA3, but not with EndoA2 (26, 29). However, specific interaction motifs have not yet been identified in these two cargoes, and it remains uncertain whether the interaction with EndoA3 is direct or indirect. If direct, it is still unknown which part of EndoA3 mediates this interaction. Nevertheless, the accumulating evidence that Ig-like cell adhesion molecules are preferentially EndoA3-dependent cargoes suggests the existence of a shared recognition pattern within this cargo family, warranting further investigations.

In addition, we found that two of the three confirmed EndoA3-dependent endocytic cargoes, ALCAM and ICAM1, are also retrograde transport cargoes dependent on the retromer complex. The fact that ICAM1 follows a retromer-dependent retrograde transport route aligns with previous proteomic analyses of retromer-dependent cargoes, where ICAM1 was identified among the hits (60). While the link between retrograde transport and cell polarity has been well-established (9–11), the relationship between CIE and subsequent retrograde transport in polarized cellular contexts has largely been unexplored. Here, we propose that EndoA3-dependent cargoes, such as ICAM1 and ALCAM, use the retrograde transport route, thereby linking this specific CIE pathway to polarized cellular environments. Interestingly, our findings align with a previous study by Jo and colleagues which suggested that ICAM1 undergoes clathrin-independent endocytosis and is transported to the immune synapse from intracellular compartments in dendritic cells (61). In their cellular model, the authors also showed that this recycling of ICAM1 was LFA1-dependent, but this remains not known and to be further explored in our model system.

The retromer complex is one of the two complexes responsible for retrieving cargoes from endosomes for retrograde transport. The other complex, named Commander, consists of the CCC and retriever complexes. The CCC complex is made up of two coiled-coil domain-containing proteins CCDC22 and CCDC93, ten COMMD (copper metabolism MURR1 (Mouse U2af1-rs1 region 1) domain) family members COMMD1-COMMD10, and DENND10 (differentially expressed in normal and neoplastic cells-containing protein 10, also called FAM45A) (7, 62–64). The retriever complex, structurally similar to the retromer complex, is composed of C16orf62 (VPS35L), DSCR3 (VPS26C), and VPS29 (6, 65). Generally, the Commander complex is considered to be responsible for retromer-independent endosomal cargo sorting for retrograde transport, although many cargoes can utilize both complexes (6). Our study confirmed that both ALCAM and ICAM1 are retromer-dependent retrograde transport cargoes. However, while depletion of the retromer subunits VPS35 and VPS26A both impaired the retrograde transport of ALCAM and ICAM1 in HeLa cells, the effect of VPS26A depletion was consistently more pronounced than that caused by the loss of VPS35 (Fig. 1C-D). Moreover, only VPS26A depletion impaired CD8 T cell activation, while VPS35 depletion had no significant effect (Fig. 3G). Western blot analyses confirmed that VPS35 siRNA treatment led to a significant reduction in VPS26A levels, whereas VPS26A siRNA treatment resulted in a milder loss of VPS35 (Fig. 1C-D, S1C and S1E). This observation is in agreement with a previous report suggesting that VPS29 and VPS35 form a stable subcomplex *in vivo*, and that depletion of VPS35 or VPS29 leads to the degradation of other subunits, whereas VPS26A depletion has minimal effects on VPS29 and VPS35 levels (66).

Based on these observations, we hypothesize that there may be a mutual compensatory relationship between the retromer and Commander complexes for some cargoes. Depletion of VPS35 in HeLa cells causes an almost complete loss of the retromer complex, potentially triggering compensation by the Commander complex. However, VPS26A depletion preserves the VPS29-VPS35 intermediate of the retromer complex (66), which may inhibit compensation by the Commander complex through mechanisms that remain to be explored. This could explain why VPS35 depletion had a milder effect on the retrograde transport of ALCAM and ICAM1 (Fig. 1C-D) and why it did not significantly affect CD8 T cell response (Fig. 3G). Future investigations on the potential mutual compensation between the retromer and Commander complexes in the retrograde transport of Ig-like proteins will be necessary.

Moreover, a striking observation from our study is that disruption of EndoA3-mediated endocytosis in cancer cells leads to the formation of larger, dysfunctional contact zones with cytotoxic CD8 T cells, accompanied by reduced cytokine secretion, yet enhanced lytic activity. This counterintuitive result aligns with established evidence that full immune synapse maturation is not required for cytotoxicity. Rather, effective killing can occur through transient and serial interactions (67). Indeed, when CD8 T cells fail to kill, they tend to form prolonged synapses and release higher levels of cytokines, whereas successful, rapid kills are associated with shorter, less stimulatory contacts (68). Single CD8 T cells, as well as CAR-T and NK cells, can kill multiple target cells sequentially through such transient contacts, so-called serial killing (69–71).

Notably, our results appear to contrast with the findings of Petit *et al.* (2016) (72), who reported that larger immune synapses correlate with increased T cell adhesion to target cells, enhanced cytokine secretion, greater degranulation, and increased lytic activity. Taken together with our data, these observations indicate that although changes in immune synapse size always reflect alterations in T cell responses, synapse size alone does not allow one to infer synapse stability or transience, nor the magnitude of T cell activation. Therefore, an in-depth characterization of T cell responses – integrating multiple readouts such as cytokine production and secretion, surface expression of activation markers, proliferation, degranulation, and cytolytic activity – is essential.

Altogether, our findings suggest that by controlling the endocytic turnover and membrane delivery of adhesion molecules such as ICAM1 at the target-cell surface, EndoA3-mediated CIE tunes the signaling properties of the immune synapse. We propose a model (Fig. 5G) where a lower EndoA3 level likely decreases ICAM1 redistribution, favoring larger but transient synapses that enhance cytolytic engagement with limited T cell cytokine secretion, which fits with the serial killing hypothesis. In contrast, active EndoA3-dependent endocytosis would promote faster adhesion-molecule turnover and formation of smaller, more stable synapses that enhance cytokine secretion by T cells, but limit their serial killing capacity.

In conclusion, we identify ICAM1 as a novel EndoA3-dependent endocytic cargo, alongside ALCAM, and show that both rely on retromer-dependent retrograde transport. Our findings connect a defined clathrin-independent endocytic mechanism to the polarized recycling of immune synapse components, positioning EndoA3-mediated endocytosis as a potential regulatory axis that fine-tunes the balance between CD8 T cell cytokine production/secretion and lytic activity. In the longer term, pharmacological modulation of this pathway may represent an interesting avenue to modulate T cell-mediated anti-tumor immunity. However, translation into therapeutic applications remains distant and will require further investigation in future studies.

## Materials and Methods

### Cell culture

Wild-type HeLa cell line was from ATCC^®^. GalT-GFP-SNAP-expressing HeLa cell line was generated in a previous study (73). EndoA3-GFP-expressing HeLa cell line was previously generated by our lab (26). HLA-A*68012-expressing HeLa cell line was generated for the current study. Wild-type LB33-MEL cell line was derived from a melanoma patient in 1988 in de Duve Institute (45). EndoA3-GFP-expressing and GalT-GFP-SNAP-expressing LB33-MEL cell lines were generated for the current study. Anti-MUM-3 CD8 T cells were derived from the same melanoma patient in 1990 in de Duve Institute (45). Epstein-Barr virus transformed B cells from cancer patient LG-2 (LG-2 EBV) were obtained as described (74).

Wild-type HeLa cells were cultured at 37°C under 5% CO_2_ in high glucose DMEM (Gibco, 41966-029) supplemented with 10% FBS (Dulis, 500105N1N), 100 U/mL penicillin and 100 µg/mL streptomycin (Penicillin-Streptomycin, Gibco, 15140-122). EndoA3-GFP-expressing, GalT-GFP-SNAP-expressing, and HLA-A*68012-expressing HeLa cells were cultured at 37°C under 5% CO_2_ in high glucose DMEM (Gibco, 41966-029) supplemented with 10% FBS, 100 U/mL penicillin, 100 µg/mL streptomycin, and 0.5 mg/mL G418 (Roche, 108321-42-2). WT LB33-MEL cells were cultured at 37°C under 5% CO_2_ in IMDM (Gibco, 12440-053) supplemented with 10% FBS, 100 U/mL penicillin, 100 µg/mL streptomycin, 1.5 mM GlutaMAX I (Gibco, 35050-038), and 0.05 mM β-mercaptoethanol (Gibco, 21985-023). Empty plasmid stably transfected (Φ), EndoA3-GFP-expressing, and GalT-GFP-SNAP-expressing LB33-MEL cells were cultured in the same medium as WT LB33-MEL cells but supplemented with 0.7 mg/mL G418 (Roche, 108321-42-2). Anti-MUM-3 CD8 T cells were freshly thawed from cryovials and cultured at 37°C under 5% CO_2_ in IMDM (Gibco, 12440-053) supplemented with 10% human serum (Ludwig Institute for Cancer Research (LICR), Brussels Branch), 100 U/mL penicillin, 100 µg/mL streptomycin, 1.5 mM GlutaMAX I, and extra 5 U/mL DNase I (STEMCELL Technologies, 07469) and 50 U/mL IL-2 (PROLEUKIN^®^, CNK 1185-958) for at least 4 h before usage. For culture, T cells were maintained with the same medium, but without DNase I. LG-2 EBV cells were cultured at 37°C under 8% CO_2_ in IMDM (Gibco, 21980-032) supplemented with 10% FBS (Gibco, A5256701), 100 U/mL penicillin, 100 µg/mL streptomycin and 1.5 mM GlutaMAX I (Gibco, 35050-038).

### DNA constructs and transfection

Plasmids used in this study are listed below:

| Protein | Backbone | Tag | Source |
| --- | --- | --- | --- |
| EndoA3 | pIRESneo2 | GFP | F. Tyckaert, LIBST, BE |
| / | pIRESneo3 | / | Clontech |
| ICAM1 | pLV-mScarlet | mScarlet | J. D. Van Buul, Amsterdam UMC, NL |
| ICAM1 | pEGFP-N1 | EGFP | Generated in the lab for this study |
| GalT-GFP-SNAP | pSNAP-tag | GFP | L. Johannes, Institut Curie, FR |
| / | pEGFP-N1 | EGFP | Clontech |
| HLA-A*68012 | pcDNA3 | / | P. van der Bruggen, de Duve Institute, BE |

C-terminally EGFP-tagged ICAM1 expression plasmid was generated in the current study for live-cell imaging. ICAM1 sequence was first amplified by PCR from the aforementioned mScarlet-tagged ICAM1 expression plasmid, and then inserted into the pEGFP-N1 vector by Gibson Assembly^®^ (New England Biolabs). To do so, the vector was linearized with the restriction enzyme EcoR I, and corresponding overlapping sequences were added to both ends of ICAM1 inserts by PCR (forward primer: 5’-CGAGCTCAAGCTTCGATGGCTCCCAGCAGC-3’; reverse primer: 5’-TACCGTCGACTGCAGGGGAGGCGTGGCTTG-3’). The Gibson assembly product was then amplified in *E. coli* DH10B strain and validated by sequencing.

For immunofluorescence and live-cell imaging experiments, plasmids were transfected with FuGENE HD (Promega), according to the manufacturer’s instructions. Cells were used for experiments 16–24 h after transfection.

To generate the stable HLA-A*68012-expressing HeLa cell line, the pcDNA3-HLA-A*68012 plasmid was transfected into wild-type HeLa cells by nucleofection (Amaxa Nucleofector 2b, Lonza) according to the built-in protocol. Transfected cells were cultured for 2 days. A selection pressure was then imposed by supplementing the culture medium with 1 mg/mL G418 for 7 days. Cells were then harvested and maintained with 0.5 mg/mL G418 in the culture medium.

For the generation of different stable LB33-MEL cell lines, 300,000 wild-type LB33-MEL cells were seeded in 2 mL culture medium. 4 µg of corresponding plasmid (pIRESneo3 for LB33-MEL Φ generation, pIRESneo2-EndoA3-GFP-FKBP for EndoA3-GFP-expressing LB33-MEL generation, and pGalT-GFP-SNAP-tag for GalT-GFP-SNAP-expressing LB33-MEL generation) were added to 200 µL of Opti-MEM (Gibco), followed by addition of 24 µL of FuGENE HD (Promega) and gentle mixing. The mix was incubated for 15 min at room temperature, and then added to the cells. After 2 days of culture, transfected cells were selected by supplementing the culture medium with 0.7 mg/mL G418 for 7-15 days. Cell sorting based on GFP fluorescence was then conducted with the FACSAria flow cytometer (BD Biosciences) to yield a 100% positive stable EndoA3-GFP-expressing LB33-MEL cell line. For LB33-MEL cells transfected with GalT-GFP-SNAP-expressing plasmid, the homogeneity of cells was checked on a FACSVerse at the end of selection, and all cells were GFP-positive. Stable LB33-MEL Φ, EndoA3-GFP-expressing LB33-MEL and GalT-GFP-SNAP-expressing LB33-MEL cell lines were then maintained with continuous presence of 0.7 mg/mL G418 in the culture medium.

### RNA interference

siRNAs were used to knock down target molecules in HeLa and LB33-MEL cell lines at a total final concentration of 40nM. For HeLa cells, 150,000 cells were seeded per well of a 6-well plate for each siRNA condition, and siRNA transfection was conducted with 1 µL Lipofectamine RNAiMAX (ThermoFisher). For LB33-MEL cells, 200,000 cells were seeded per well of a 6-well plate for each siRNA condition, and siRNA transfection was conducted with 2 µL Lipofectamine RNAiMAX (ThermoFisher). All experiments were done 72 h after siRNA transfection. All siRNAs were purchased from QIAGEN, and their sequences are listed as follows:

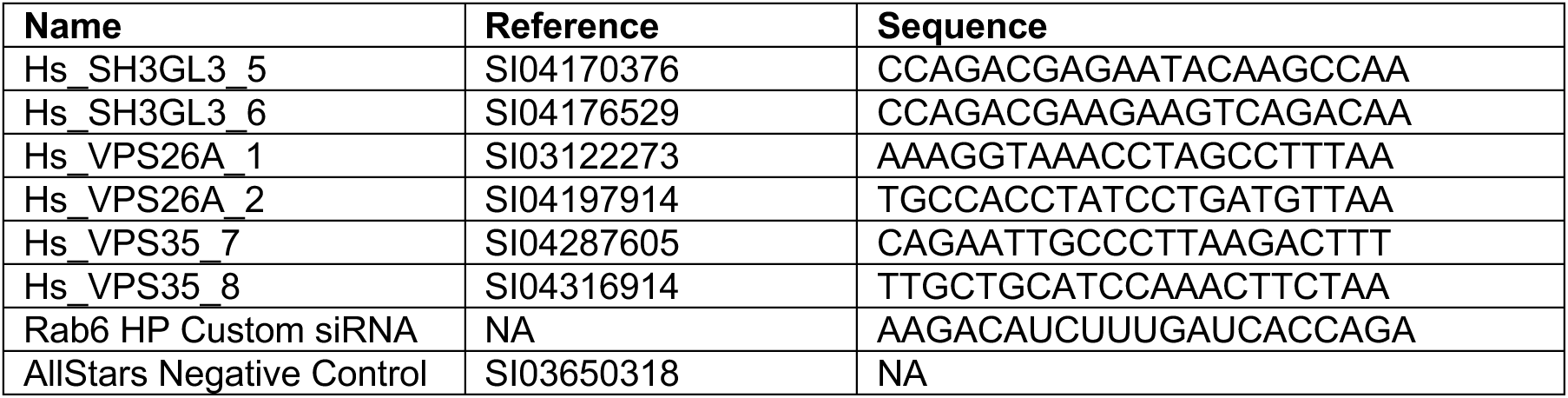

### T cell proliferation assays and quantification of activation markers

For data of Fig. S4G-J, siRNA-treated LB33-MEL EndoA3+ cells were detached with PBS-EDTA 2 mM, resuspended to 300,000 cells/mL, and γ-irradiated (100 Gy), as were LG2-EBV feeder cells. Anti-MUM-3 CD8 T cells were thawed the day before, labeled with 1 µM CellTrace Violet (Invitrogen™, C34571) for 20 min at 37°C, washed, and resuspended to 300,000 cells/mL in T cell medium supplemented with 50 U/mL IL-2. In ultra-low attachment round-bottom 96-well plates (Corning® Costar®, CLS7007), 30,000 CD8 T cells were co-seeded with 15,000 LB33-MEL cells and 90,000 feeder cells. Plates were briefly centrifuged and incubated for 4 days at 37°C with 8% CO₂. Cells were then placed on ice for antibody staining and flow cytometry analysis.

For data of Fig. S5B, HLA-A*68012 expressing HeLa cells were detached with PBS-EDTA 2 mM and resuspended to 1,000,000 cells/mL in the culture medium supplemented with or without MUM-3 peptide (EAFIQPITR) at 300 ng/mL. Then, HeLa cells and LG-2 EBV feeder cells were γ-irradiated for 45 min (100 Gy), washed, and maintained in T cell culture medium until the initiation of co-culture with T cells. Anti-MUM-3 CD8 T cells were thawed the day before and were harvested, washed, and resuspended in complete T cell culture medium with 0.5 µM CMFDA (Invitrogen™, C2925) for 20 min at 37°C. T cells were then washed and resuspended to 300,000 cells/mL in T cell medium supplemented with 50 U/mL IL-2. In an ultra-low attachment round bottom 96-well plate (Corning^®^ Costar^®^, CLS7007), 30,000 anti-MUM-3 CD8 T cells were co-seeded with 10,000 HeLa cells and 90,000 γ-irradiated LG-2 EBV feeder cells. The plate was quickly centrifuged and then incubated for 4 days at 37°C with 8% CO_2_. Cells were then transferred on ice for antibody staining and flow cytometry analysis.

Flow cytometry data were acquired with a LSR Fortessa (BD Biosciences) and analyzed using FlowJo (v10.8.1) software.

### T cell IFNγ secretion assay

CD8 T cells were thawed and cultured as described above. siRNA transfected LB33-MEL cells were harvested 72 h after the siRNA transfection and were resuspended to 300,000 cells/mL in T cell medium. CD8 T cells were then washed and resuspended to 300,000 cells/mL in fresh T cell medium supplemented with 50 U/mL IL-2. 100 µL of LB33-MEL cells and 100 µL of T cells were pooled together in a 96-well plate and were co-cultured for 20 h at 37°C under 5% CO_2_. A technical triplicate was made for each siRNA condition. The supernatant was collected the next day for ELISA analysis.

ELISA plate was coated with 50 µL IFNγ monoclonal coating antibody solution (Invitrogen, AHC4432; 1:250 dilution in PBS) in each well and was incubated at 4°C overnight. On the next day, two standard IFNγ concentration ladders were prepared: 1:2 dilutions for 7 successive times with T cell medium in 96-well plate (original concentration of 10,000 pg/mL; the last well contains only medium). The ELISA plate was washed with washing buffer (0.154 M NaCl solution with 0.1% Tween-20) twice and with water once to remove unbound antibody. 50 µL of supernatant and standard IFNγ ladders were then added to the ELISA plate, followed by the addition of 50 µL IFNγ monoclonal detecting antibody solution (biotin conjugated; Invitrogen, AHC4539; 1:1000 dilution in T cell medium) into each well. The plate was incubated on a shaker at 37°C for 2 h, and then was washed again. 50 µL of Streptavidin-Peroxydase solution (Sigma, S5512; 1:1000 dilution in PBS containing 0.5% bovine serum albumin) was added to each well and the plate was again incubated at 37°C for 30 min. Then, the plate was washed and 100 µL of 1-Step Ultra TMB-ELISA Substrate Solution (ThermoFisher, 34028) was added to each well. When the blue color of TMB stopped getting darker, 20 µL of sulfuric acid (1 M) was added to each well to stop the reaction (the blue color turned yellow). The absorbance of substrate solution at 450 nm was then measured with an ELISA Analyzer (Bio-Rad Microplate Reader) and the IFNγ concentration was determined.

### Quantification of intracellular cytokines in CD8 T cells

CD8 T cells were thawed and cultured as described above. LB33-MEL cells were harvested 72 h after siRNA transfection and resuspended to 1,000,000 cells/mL in T cell medium. T cells were then washed and resuspended to 1,000,000 cells/mL in fresh T cell medium supplemented with 50 U/mL IL-2. 100 µL LB33-MEL cells and 100 µL T cells were pooled together in a 96-well “U” bottom plate and were co-cultured at 37°C under 8% CO_2_ for 1 h. Then, Brefeldin A (BFA; Sigma, B7651-5MG) was added to each well at a final concentration of 5 µg/mL. Co-culture continued for another 3 h, after which cells were transferred to a 96-well “V” bottom plate and washed with 150 µL/well of staining buffer (PBS with 1mM EDTA and 1% human serum). The plate was quickly spun at 400×g for 3 min and supernatant was removed. 50 µL of extracellular staining antibody solution for CD8 and viability marker prepared with staining buffer were added to each well and mixed. The plate was then maintained at 4°C for 20 min in the dark. Afterwards, cells were washed with 150 µL/well PBS and resuspended in 100 µL/well of 1% paraformaldehyde (PFA) for 10 min at room temperature. Cells were then washed with PBS again and resuspended in 100 µL PBS containing 1 mM EDTA, 1% human serum, and 0.1% saponin. After 10 min incubation at room temperature, cells were centrifuged, and supernatant was removed. 50 µL of intracellular staining antibody solution for cytokines prepared with the same buffer was added to each well, and the plate was incubated in the dark for 30 min at room temperature. Finally, cells were washed and resuspended in 250 µL 1% PFA for flow cytometry analysis.

Flow cytometry data were acquired with a FACSVerse (BD Biosciences) and were analyzed using FlowJo (v10.8.1) software.

### T cell degranulation assay

CD8 T cells were thawed and cultured as described above. LB33-MEL cells were harvested 72 h after siRNA transfection and resuspended to 1,000,000 cells/mL in T cell medium. T cells were then washed and resuspended to 2,000,000 cells/mL in fresh T cell medium supplemented with 50 U/mL IL-2. 100 µL LB33-MEL cells and 50 µL T cells were pooled together in a 96-well “U” bottom plate with 10 µL anti-CD107a antibody. Cells were co-cultured at 37°C under 8% CO_2_ for 10 min, and were then transferred on ice for extracellular marker and viability dye staining and flow cytometry analysis.

### T cell cytotoxicity assay

Anti-MUM-3 CD8 T cells were thawed one day before the experiment. siRNA-transfected LB33-MEL EndoA3+ cells were harvested 72 h post-transfection and resuspended to 1×10⁶ cells/10 µL FBS. Cells were labeled with 20 µL Chromium-51 (PerkinElmer, NEZ030S002MC) for 1 h at 37°C. After two centrifugations (400xg for 8 min), cells were resuspended in 10 mL medium and incubated for an additional hour. After washing, labeled cells were counted and adjusted to 10,000 cells/mL.

CD8 T cells were washed and resuspended to 300,000 cells/mL. In a 96-well V-bottom plate, 150 µL of T cells were dispensed and a 3-fold serial dilution was performed. Then, 100 µL labeled LB33-MEL cells were added, and plates were incubated for 4 h at 37°C with 8% CO₂. Supernatants (100 µL) were mixed with 150 µL scintillation fluid (PerkinElmer, 6013117) in a chromium-reading plate (PerkinElmer, 1450-40). Radioactivity was measured on a Microbeta2 counter (PerkinElmer). Technical triplicates were included for each siRNA condition.

### Cell surface staining of cancer cells for flow cytometry

72 h after siRNA transfection, cells were detached and harvested with PBS-EDTA 4 mM and were transferred to pre-cooled 96-well “V”-bottom plates with 100,000 cells in each well. The plate was centrifuged at 400xg for 4 min at 4°C. Cells were washed with 150 µL/well of staining buffer (PBS with 1mM EDTA and 1% human serum) and centrifuged at 400xg for 4 min at 4°C. Cells were resuspended in 50μL extracellular staining antibody solution for ALCAM and ICAM1 and viability marker, prepared with staining buffer. The plate was then maintained at 4°C for 20 min in darkness. Finally, cells were washed and resuspended in 250 µL 1% PFA for flow cytometry analysis.

Flow cytometry data were acquired with a FACSVerse (BD Biosciences) and were analyzed using FlowJo (v10.8.1) software.

### Western blotting

Cell lysates were prepared with 200,000 cells and 50 µL of sample buffer (62.5 mM Tris-HCl pH 6.8, 2% sodium dodecyl sulfate (SDS), 5% glycerol, 2.5% β-mercaptoethanol, 0.005% blue bromophenol). Lysate samples were boiled at 95°C for 10 min before loading onto 4–15% Mini-PROTEAN^®^ TGX™ precast protein gels (Bio-Rad, 4561084), as well as prestained protein standard. For the SNAP-Tag-based retrograde transport assay (see below), HiMark™ Pre-stained Protein Standard (Invitrogen, LC5699) was used; for all the other western blots, Broad Range (10-250 kDa) Color Prestained Protein Standard (New England Biolabs, P7719S) was used. After electrophoresis, proteins were transferred onto a PVDF membrane (pre-activated with methanol for 1 min; Millipore, IPFL85R) using a wet transferring system. The membrane was then blocked in 5%-milk PBS-T (phosphate-buffered saline, 0.1% Tween-20) for 1h at room temperature, then stirred overnight at 4°C with primary antibody in the same solution. The next day, the membrane was incubated with corresponding horseradish peroxidase (HRP) conjugated secondary antibodies for 1h at room temperature. Chemiluminescent (ECL) substrate (SuperSignal™ West Femto Maximum Sensitivity Substrate, ThermoFisher, 34096, or SuperSignal™ West Pico PLUS Chemiluminescent Substrate, ThermoFisher, 34580) was used for image exposure with an Amersham ImageQuant™ 800 machine (Cytiva). Images were processed and quantified with ImageJ v2.14.0/1.54f (NIH) software.

### SNAP-Tag-based retrograde transport assay

First, anti-ALCAM (Bio-Rad, MCA1926) and anti-ICAM1 (Bio-Rad, MCA1615) antibodies were labeled with benzylguanine (BG) by incubating them overnight at 4°C with a threefold molar excess of BG-GLA-NHS reagent (New England Biolabs, S9151S; prepared in anhydrous DMSO). The reaction was neutralized with 20 mM Tris (pH 7.4) for 10 min at 21°C. Free BG-GLA-NHS was eliminated by applying labelled antibodies on spin desalting columns with a 7 kDa cutoff (ThermoFisher, 89849), according to manufacturer’s instructions. Then, retrograde transport assays were conducted according to previously published protocols (44).

### Colocalization and uptake experiments

LB33-MEL cells were cultured on coverslips in 4-well plates to reach sub-confluence on the day of the experiment. Cells were pre-incubated in fresh serum-containing medium for 30 min at 37°C just before the experiment, then were incubated with anti-ALCAM antibody (Biolegend, 343902)-containing pre-warmed medium (5 µg/mL) for 15 min at 37°C to allow for endocytosis. For colocalization experiments, cells were quickly washed at 37°C with pre-warmed PBS and were subsequently fixed with pre-warmed 4% PFA for 20 min at 37°C in order to preserve the integrity of cellular structures and protein distribution. For uptake experiments, endocytosis was stopped on ice, and unbound antibodies were removed by extensive washes with ice-cold PBS++ (PBS supplemented with 0.4 mM MgCl_2_ and 0.9 mM CaCl_2_). Residual cell surface-accessible anti-ALCAM antibodies were stripped by two 5 s acid washes on ice (200 mM acetic acid, pH 2.5, 300 mM NaCl, 5 mM NaCl, 1 mM CaCl_2_, and 1 mM MgCl_2_). After neutralization by extensive washes with ice-cold PBS++, cells were fixed with 4% PFA for 10 min on ice, followed by another 10 min at room temperature. For both experiments, after quenching with 50 mM NH_4_Cl for at least 5 min, cells were permeabilized with 0.02% saponin in PBS containing 0.2% bovine serum albumin for 30 min at room temperature. After incubation with secondary antibodies in the same permeabilization solution for 30 min, samples were mounted with Fluoromount G (Invitrogen). Images were taken with a LSM900 microscope (Carl Zeiss) equipped with an Airyscan detector and a Plan Apo 63× numerical aperture (NA) 1.4 oil immersion objective. For colocalization experiments, Airyscan mode was applied. Pixel size: 0.04 micron. For uptake experiments, confocal mode was used. Pixel size: 0.07 micron. Microscope software: Zen blue (v3.3; Carl Zeiss).

### Conjugate formation for confocal fixed sample imaging

For conjugate formation between adherent HLA-A*68012-expressing HeLa cells and anti-MUM-3 CD8 T cells (Fig. S5C), HeLa cells were cultured on coverslips and transiently transfected with ICAM1-mScarlet expression plasmid 16-24h before the experiment. HeLa cells were first incubated in 300 ng/mL MUM-3 peptide-containing culture medium for 45 min at 37°C and were then washed extensively with T cell culture medium to remove residual unbound peptide. T cells were freshly thawed as described above and resuspended to 500,000 cells/mL with T cell culture medium supplemented with 50 U/mL IL-2. Then, 250,000 T cells were added to HeLa cells and the pool was incubated for 20 min at 37°C. Cells were then fixed at room temperature for 20 min by adding PFA directly to the culture medium to reach a final concentration of 4%.

For conjugate formation between suspended HLA-A*68012-expressing HeLa cells and anti-MUM-3 CD8 T cells (Fig. 5A, S5D), HeLa cells were cultured in 6-well plates and transfected with negative control siRNAs or siRNAs targeting EndoA3 72h in advance and with plasmid expressing ICAM1-mScarlet 16-24h in advance. HeLa cells were detached with PBS-EDTA 4 mM, resuspended to 1,000,000 cells/mL in culture medium containing 300 ng/mL MUM-3 peptide and incubated for 45 min at room temperature. Then, HeLa cells were extensively washed with T cell culture medium, and finally resuspended to 1,000,000 cells/mL with T cell culture medium supplemented with 50 U/mL IL-2. T cells were freshly thawed as described above and resuspended to 1,000,000 cells/mL with T cell culture medium supplemented with 50 U/mL IL-2. Subsequently, 250,000 HeLa cells and 250,000 T cells were pooled together and further co-cultured on coverslips for 20 min at 37°C. Of note, coverslips were pre-coated with poly-L-lysine (Sigma-Aldrich, P4832) according to manufacturer’s instructions to stabilize conjugates. Cells were then fixed at room temperature for 20 min by adding PFA directly to the culture medium to reach a final concentration of 4%.

For conjugate formation between suspended LB33-MEL EndoA3+ cells and anti-MUM-3 CD8 T cells (Fig. 5C and 5E, S6A-B), LB33-MEL cells were transfected with negative control siRNAs or siRNAs targeting EndoA3 or VPS26A 72h in advance. LB33-MEL cells were detached with 0.05% Trypsin-EDTA (Gibco, 25300-054) and resuspended to 1,000,000 cells/mL with T cell culture medium supplemented with 50 U/mL IL-2. T cells were freshly thawed as described above and resuspended to 1,000,000 cells/mL with T cell culture medium supplemented with 50 U/mL IL-2. Then, 250,000 LB33-MEL cells and 250,000 T cells were pooled together and were co-cultured on poly-L-lysine pre-coated coverslips for 20 min at 37°C. Cells were then fixed at room temperature for 20 min by adding PFA directly to the culture medium to reach a final concentration of 4%.

After fixation with PFA and quenching with 50 mM NH_4_Cl for at least 5 min, cells were permeabilized with 0.02% saponin in PBS containing 0.2% bovine serum albumin for 30 min, both at room temperature. Then, cells were successively incubated with primary and secondary antibodies in the same PBS-saponin solution for 30 min and mounted with Fluoromount G (Invitrogen). Images were taken with LSM900 microscope (Carl Zeiss) equipped with an Airyscan detector and a Plan Apo 63× numerical aperture (NA) 1.4 oil immersion objective. For the ICAM1 recruitment experiment (Fig. S5C) and immune synapse morphology experiment (Fig. 5C and 4E, S6A-B), z-stack was used. z-stack interval: 0.15 micron. For all images, Airyscan mode was applied. Pixel size: 0.04 micron. Microscope software: Zen blue (v3.3; Carl Zeiss).

### Conjugate formation for high-speed spinning-disk confocal live-cell imaging

HLA-A*68012-expressing HeLa cells were seeded in 8-well culture chamber slides (Ibidi, 80806) and transfected with ICAM1-mScarlet expression plasmid 16-24 h in advance. HeLa cells were first incubated in MUM-3 peptide-containing (300 ng/mL) culture medium for 45 min at 37°C and then washed extensively with T cell culture medium to remove residual unbound peptide. Anti-MUM-3 CD8 T cells were freshly thawed as described above and stained with CellTracker Green CMFDA (Invitrogen, C2925; 1:100 dilution in T cell culture medium supplemented with 50 U/mL IL-2) for 45 min at 37°C, washed with culture medium, and resuspended to 300,000 cells/mL in T cell culture medium supplemented with 50 U/mL IL-2. 100,000 T cells were added to each well containing HeLa cells and live-cell imaging was started immediately after. Live-cell imaging was performed at 5% CO_2_ and 37°C on an inverted Axio Observer 7 microscope (Carl Zeiss) equipped with a Yokogawa CSU-W1 spinning-disk confocal scanner unit, a sCMOS Photometrics Prime 95B camera, an α-Plan Apo 100× numerical aperture (NA) 1.46 oil immersion objective and a stage-top incubator. The microscope was controlled by Metamorph software (v7.10.4, Molecular Devices). Focus was maintained with Definite Focus 3 (Carl Zeiss). Several cells were imaged each for 3 minutes, with a time interval between each frame of 1 s. Pixel size: 0.11 micron.

### TIRF live-cell imaging

EndoA3-GFP-expressing HeLa/LB33-MEL cells were cultured on a 35 mm imaging dish with a glass bottom and low walls (ibidi, µ-Dish 35 mm, low, 80137) and transfected with ICAM1-mScarlet expression plasmid 16-24 h in advance. Live-cell imaging was conducted under 5% CO_2_ and 37°C with Axio Observer 7 microscope (Carl Zeiss) equipped with an iLas azimuthal TIRF module (Gattaca Systems), an α Plan-Apo 100× NA 1.46 oil immersion objective, an sCMOS Prime BSI Express camera (Photometrics) and a stage-top incubator (Tokai Hit). Microscope was controlled by Metamorph^®^ software (v7.10.4.407; Molecular Devices). Laser focus was maintained with definite focus 3. Time interval between each frame: 1 s. Pixel size: 0.065 micron.

### Cancer cell area, aspect ratio and roundness analysis by confocal imaging

For quantification of area, aspect ratio and roundness of LB33-MEL EndoA3+ cells following EndoA3 depletion, cells were seeded on coverslips and transfected with siRNAs for 72 h. Cells were then fixed with 4% PFA for 20 min and a quenching step with 50 mM NH_4_Cl was performed for at least 5 min. Cells were subsequently incubated with DAPI (Roche, 10236276001) and Phalloidin-Alexa Fluor 633 (Invitrogen, A22284, 1:500 dilution in a PBS solution containing 0.02% saponin and 0.2% bovine serum albumin) for 1 h at room temperature. Finally, coverslips were mounted with Fluoromount G (Invitrogen). Images were acquired on a Cell Discoverer 7 (Carl Zeiss) in confocal mode, using a Plan Apo 5x numerical aperture (NA) 0.35 objective and a 2x tube lens. The microscope was controlled by Zen Blue (v3.7.97.09, Carl Zeiss). Pixel size: 0.282 micron.

### Microscope image analyses and quantifications

Airyscan image processing was performed with Zen blue (v3.3; Carl Zeiss). Images were quantified with (Fiji Is Just) ImageJ v2.14.0/1.54f (NIH) software as follows.

For CD166 uptake experiments, images were quantified using Icy v2.4.2.0 (Institut Pasteur) software. Briefly, regions of interest (ROIs) were drawn manually around each cell. Within each ROI, bright spots corresponding to endocytic structures were automatically detected over dark background using the “Spot Detector” plugin. The plugin performs image denoising by computing wavelet adaptive threshold (WAT) on the union of all ROIs present in an image. This automatic thresholding was then manually adjusted, depending on the size of the spots. For each independent experiment, scale 2 was chosen and sensitivity was adjusted empirically. In addition, a size filter was added to discard spots smaller than 2 pixels. After processing, an excel file containing the fluorescence intensity of each endocytic structure was generated. The sum intensity of endocytic structures in each cell was calculated, then data were first adjusted according to the surface level of CD166 detected by flow cytometry in each experimental condition and were finally normalized to the control condition (set as 100%) for statistical analysis.

For conjugate formation experiments with HeLa cells to detect ICAM1 recruitment in the vicinity of CD8 T cells, masks were automatically generated for each image according to phalloidin staining by using Cellpose v3.0.7 software (75). First, an RGB format 512 × 512-pixel image of the phalloidin channel was generated from Fiji (ImageJ v2.14.0/1.54f) for each image and was imported to Cellpose. Segmentation of each cell was automatically performed using “cyto3” model with diameter (pixels) set as 40 for all independent repeats and was then exported as a mask. The mask was then imported into Fiji and re-scaled to the real size of the original image. Wand tool was used to select ROI corresponding to single cells. In each ROI, the raw integrated density (RawIntDen) of ICAM1 signal was measured. Then, the ICAM1 recruitment ratio in the vicinity of the T cell was calculated as the ratio between the ICAM1 signal in the T cell ROI and the total ICAM1 signal in the ROIs of the T cell and the adjacent HeLa cell: “RawIntDen of ICAM1 signal in T cell ROI/(RawIntDen of ICAM1 signal in T cell ROI + RawIntDen of ICAM1 signal in HeLa cell ROI)”. Data were normalized to control condition (set as 100%) for each independent repeat and were then pooled together for statistical analysis.

For conjugate formation experiments to measure the size of immune synapses, z-stack images (z-stack interval: 0.15 micron) for each conjugate were imported to Fiji. The contact region between a CD8 T cell and an LB33-MEL cell was delineated by “freehand line” tool manually, and then the length of the line was measured for each z-stack. The largest value was picked out from all z-stacks and was considered as the size of this immune synapse. All data from three independent repeats were then pooled together for statistical analysis.

Tracking of ICAM1-positive carriers in live-cell spinning-disk images of HeLa cells upon formation of immune synapses with CD8 T cells was performed using Fiji software (ImageJ v2.14.0/1.54f). Segmentation of moving carriers was first carried out by machine learning using the Trainable Weka Segmentation plugin (76). Tracking was then performed with TrackMate (77), using the Advanced Kalman Tracker. Resulting tracks were filtered based on their duration, displacement and quality to refine the dataset and remove non-moving structures. Track counts within defined cellular regions were calculated and normalized to the surface area of each region to obtain track densities. Track displacements and directions (for polar plot) were computed separately in Python using a custom script.

For quantification of area, aspect ratio and roundness of LB33-MEL EndoA3+ cells upon EndoA3 depletion, Arivis software (v4.1.1, Carl Zeiss) was used. Cells were segmented using Cellpose (78) Cyto2 model based on phalloidin and DAPI staining.

### Statistical analyses

Statistical analyses were performed with GraphPad Prism v10.1.0. D’Agostino–Pearson omnibus normality test was used to check the normality of datasets. If data passed the normality test, parametric tests were used, and data were plotted on graphs as mean ± SEM as error bars. If data did not pass the normality test, nonparametric tests were used, and data were plotted on graphs as median and quartiles. Details on the parametric and nonparametric tests used for each analysis are indicated in figure legends. Significance of comparison is represented on the graphs by asterisks.

### Antibodies and reagents

| Product | Reference | Application | Dilution |
| --- | --- | --- | --- |
| PE anti-human CD166 antibody | Biolegend, 343904 | FACS | 1:1000 |
| APC anti-human CD54 antibody | BD Pharmingen, 561899 | FACS | 1:60 |
| APC anti-HLA-A2, A28 antibody | Miltenyi Biotec, 130-099-582 | FACS | 1:300 |
| Viability Dye eFluor 780 | Invitrogen, 65-0865-14 | FACS | 1:1000 |
| PerCP-Cy5.5 anti-human CD8 antibody | BD Pharmingen, 624060 | FACS | 1:100 |
| APC anti-human IL-2 antibody | Biolegend, 500310 | FACS | 1:25 |
| BV421 anti-human TNF-α antibody | Biolegend, 502932 | FACS | 1:100 |
| PE anti-human IFNγ antibody | BD Pharmingen, 554701 | FACS | 1:200 |
| Biotin anti-human HLA-A2 Antibody | Biolegend, 343322 | FACS | 1:400 |
| PE-Cy7 anti-human CD107a | Biolegend, 328618 | FACS | 1:20 |
| BV785 anti-human PD-1 antibody | Biolegend, 329930 | FACS | 1:40 |
| PE anti-human CD137 antibody | Biolegend, 309804 | FACS | 1:200 |
| PE-Cy7 anti-human TIM-3 | Biolegend, 345014 | FACS | 1:50 |
| CMFDA | Invitrogen, C2925 | FACS | 0.5 μM |
| Anti-IFNγ coating antibody | Invitrogen, AHC4432 | ELISA | 1:250 |
| Biotin anti-IFNγ detecting antibody | Invitrogen, AHC4539 | ELISA | 1:1000 |
| Streptavidin–Peroxidase | Sigma-Aldrich, S5512 | ELISA | 1:1000 |
| 1-Step Ultra TMB-ELISA Substrate | ThermoFisher, 34028 | ELISA | 100 μL |
| Anti-SH3GL3/Endophilin-A3 antibody | Sigma-Aldrich, HPA039381 | WB | 1:1000 |
| Anti-α-tubulin antibody | Sigma-Aldrich, T5168 | WB | 1:10000 |
| Anti-VPS26A antibody | abcam, ab23892 | WB | 1:1000 |
| Anti-VPS35 antibody | abcam, ab10099 | WB | 1:500 |
| HRP conjugated anti-mouse antibody | Agilent Technologies, P044701-2 | WB | 1:5000 |
| HRP conjugated anti-rabbit antibody | Agilent Technologies, P044801-2 | WB | 1:5000 |
| HRP conjugated anti-goat antibody | Santa Cruz Biotechnology, sc-2354 | WB | 1:1000 |
| Anti-SNAP-tag<sup>®</sup> antibody | New England Biolabs, P9310S | WB | 1:1000 |
| Anti-Rab6 antibody | Abcam, ab95954 | WB | 1:1000 |
| Anti-CD166 antibody | Bio-Rad, MCA1926 | SNAP-Tag | 10 μg/mL |
| Anti-CD54 antibody | Bio-Rad, MCA1615 | SNAP-Tag | 10 μg/mL |
| SNAP-Cell<sup>®</sup> Block | New England Biolabs, S9106S | SNAP-Tag | 1:200 |
| Anti-CD166 antibody | Biolegend, 343902 | IF | 1:100 |
| Anti-CD45 antibody | Santa Cruz Biotechnology, sc-1178 | IF | 1:100 |
| SNAP-Cell<sup>®</sup> TMR-Star | New England Biolabs, S9105S | IF | 1:200 |
| Anti-GM130 antibody | Biolegend, 937002 | IF | 1:200 |
| Alexa Fluor<sup>™</sup> 488 Phalloidin | Invitrogen, A12379 | IF | 1:200 |
| Alexa Fluor<sup>™</sup> 568 Phalloidin | Invitrogen, A12380 | IF | 1:200 |
| Alexa Fluor<sup>™</sup> 488 anti-Rabbit IgG | Invitrogen, A11008 | IF | 1:200 |
| Alexa Fluor<sup>™</sup> 568 anti-Mouse IgG | Invitrogen, A11031 | IF | 1:200 |
| Alexa Fluor™ 633 anti-Mouse IgG | Invitrogen, A21050 | IF | 1:200 |
| CellMask™ Plasma Membrane Stain | Invitrogen, C10045 | IF | 1:1000 |
| DAPI | Sigma-Aldrich, D9542 | IF | 1:10000 |
| Poly-L-lysine | Sigma-Aldrich, P4832 | IF | NA |

## Acknowledgments

We thank J.D. van Buul (Amsterdam UMC, Netherlands) for kindly providing aforementioned plasmid. We thank E. Rigaux, F. Tyckaert, C. Wildmann, C. Duhamel, S. Meurant, A. Fattaccioli, S. Burteau and C. Demazy for their support and technical help in experiments. We thank E. Macdonald for proofreading the manuscript. We greatly thank Morph-Im platform of UNamur and flow cytometry & cell sorting platform (CYTF) from de Duve Institute of UCLouvain for giving access to advanced microscopy and flow cytometry technologies, respectively.

S.X. is supported by a PhD fellowship from FSR (Fond Spécial de Recherche) of UNamur and UCLouvain. A.B. is a FRIA (Fonds pour la Formation à la Recherche dans l’Industrie et l’Agriculture) PhD Research Fellow from the Fonds de la Recherche Scientifique (FNRS, Belgium). T.H. is a postdoctoral research fellow of the FNRS (Belgium). L.T. is supported by a Marie Skłodowska-Curie postdoctoral fellowship (grant agreement n° 101151524) under the European Union’s Horizon 2020 program, and by an Honorary Postdoctoral Research Fellowship of the FNRS (Belgium) and an additional operating grant from the King Baudouin Foundation (Belgium). L.J. is supported by Mizutani Foundation for Glycosciences (reference n° 200014), Agence National de la Recherche (ANR-20-CE15-0009-01, ANR-22-CE11-0030-03, France), Fondation pour la Recherche Médicale (EQU202103012926, France) and an ERC Proof of Concept (project 101062030). P.V.D.B. and T.H. were supported by de Duve Institute (Belgium) and Université Catholique de Louvain (Belgium). P.M. is supported by a PDR (T.0163.21) and CDR (J.0127.23) from the Fonds de la Recherche Scientifique (FNRS, Belgium). H.-F.R. is supported by a Start-Up Grant Collen-Francqui from Francqui Foundation (Belgium), an Incentive Grant for Scientific Research (MIS-F.4540.21) and a Research Credit (CDR-J.0176.24) from the Fonds de la Recherche Scientifique (FNRS, Belgium), and a research grant from NARILIS institute (UNamur).

## Supporting Information

**Fig. S1.**
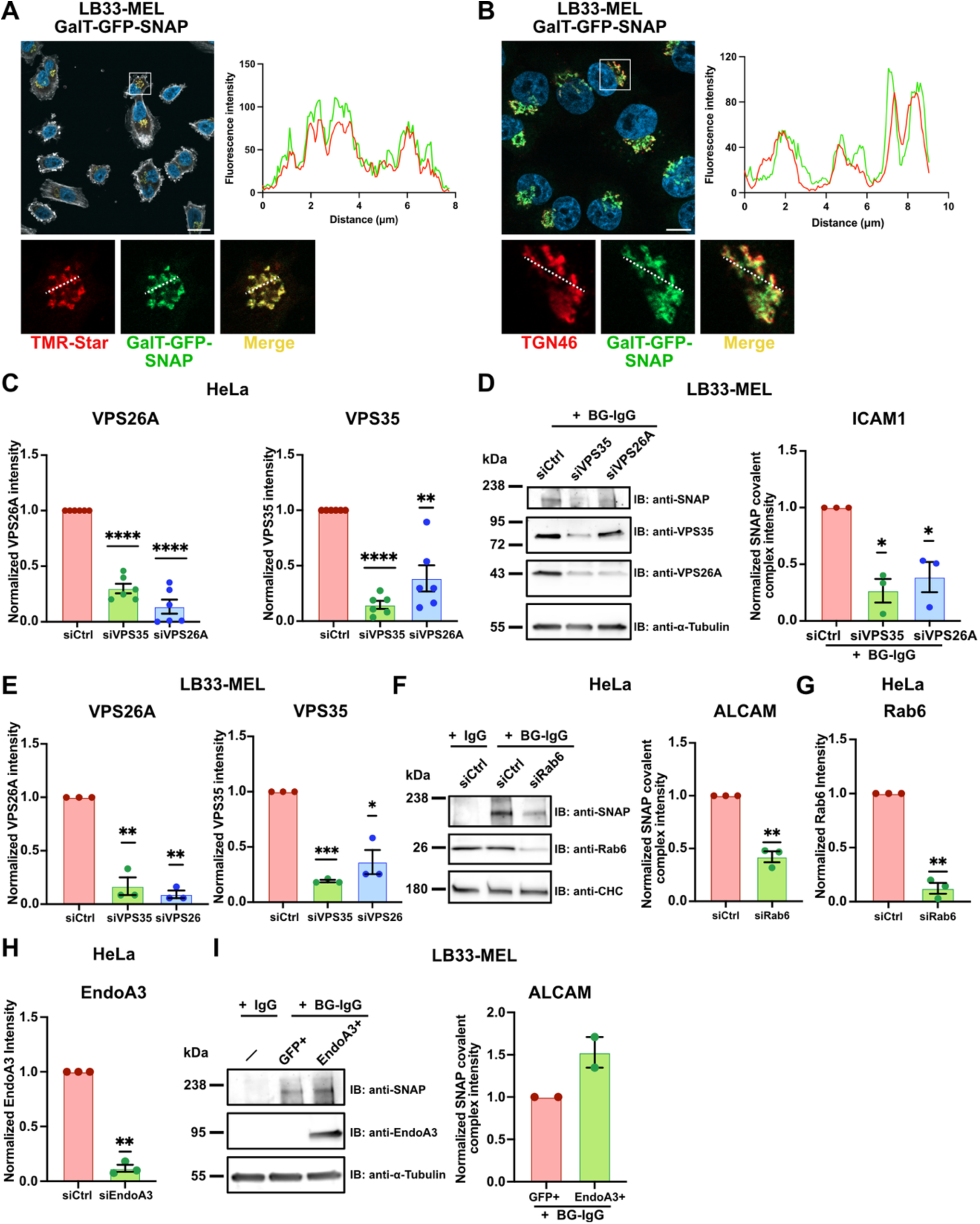
Immune synapse components ALCAM and ICAM1 are retrograde transport cargoes that rely on EndoA3, retromer and Rab6. A Confocal images of GalT-GFP-SNAP (green) and TMR-Star (red) in LB33-MEL cells stably expressing the Golgi-resident GFP-fused SNAP-tag construct (LB33-MEL GalT-GFP-SNAP). Actin (phalloidin, white) and nuclei (DAPI, blue) were also stained. The fluorescence intensity profile was made along the dashed line region in enlarged cropped area, and shows the colocalization of both signals. Scale bar: 20 μm. B Confocal images of GalT-GFP-SNAP (green) and TGN46 (red) in LB33-MEL GalT-GFP-SNAP cells. Nuclei (DAPI, blue) were also stained. The fluorescence intensity profile was made along the dashed line region in enlarged cropped area, and shows the colocalization of both signals, indicating that GalT-GFP-SNAP is correctly localized at the *TGN*. Scale bar: 10 μm. C Quantifications of the immunoblots shown in Fig. 1C-D (IB: anti-VPS35 and anti-VPS26A) confirm depletion efficiency of VPS35 and VPS26A in HeLa GalT-GFP-SNAP cells. D Retrograde transport of ICAM-1. Continuous BG-labelled anti-ICAM1 antibody uptake for 4h at 37°C in LB33-MEL GalT-GFP-SNAP cells. Western blot analysis of LB33-MEL GalT-GFP-SNAP cells transfected with siRNAs: negative control (siCtrl) or against retromer subunits (siVPS35 and siVPS26A). Immunodetection with anti-SNAP, anti-VPS35, anti-VPS26A and anti-α-Tubulin (loading control) antibodies. Quantification of the covalent IgG-SNAP complex is shown as fractions of siCtrl condition (histogram). Quantification of VPS35 and VPS26A depletion is shown in Fig. S1E. E Quantifications of the immunoblots shown in Fig. S1D confirm depletion efficiency of VPS35 and VPS26A in LB33-MEL GalT-GFP-SNAP cells. F Retrograde transport of ALCAM. Continuous BG-labelled anti-ALCAM antibody uptake for 4h at 37°C in HeLa GalT-GFP-SNAP cells. Western blot analysis of HeLa GalT-GFP-SNAP cells transfected for 72h with siRNAs: negative control (siCtrl) or against Rab6 (siRab6). Immunodetection made with anti-SNAP, anti-Rab6 and anti-clathrin heavy chain (CHC, loading control) antibodies. Quantification of the covalent IgG-SNAP-GFP-GalT complex is shown as fractions of siCtrl condition (histogram). Quantification of Rab6 depletion is shown in Fig. S1G. G Quantifications of the immunoblots shown in Fig. S1F confirm depletion efficiency of Rab6 in HeLa GalT-GFP-SNAP cells. H Quantifications of the immunoblots shown in Fig. 1E-F (IB: anti-EndoA3) confirm depletion efficiency of EndoA3 in HeLa GalT-GFP-SNAP cells. I Retrograde transport of ALCAM. Continuous BG-labelled anti-ALCAM antibody uptake for 4h at 37°C in LB33-MEL GalT-GFP-SNAP cells. Western blot analysis of LB33-MEL GalT-GFP-SNAP cells transfected (or not) with plasmids encoding free GFP (GFP+) or EndoA3-GFP (EndoA3+). Immunodetection with anti-SNAP, anti-EndoA3 and anti-α-Tubulin (loading control) antibodies. Quantification of the covalent IgG-SNAP-GFP-GalT complex is shown as fractions of the GFP+ condition (histogram). Data information: In (A-B), images are from a single experiment. In (C), data are pooled from six independent experiments. In (D-H), data are pooled from three independent experiments. In (I), data are pooled from two independent experiments. Data are presented as mean ± SEM. *P<0.05, **P < 0.01, ***P < 0.001, ****P < 0.0001. One sample *t* and Wilcoxon test. Source data are available online for this figure.

**Fig. S2.**
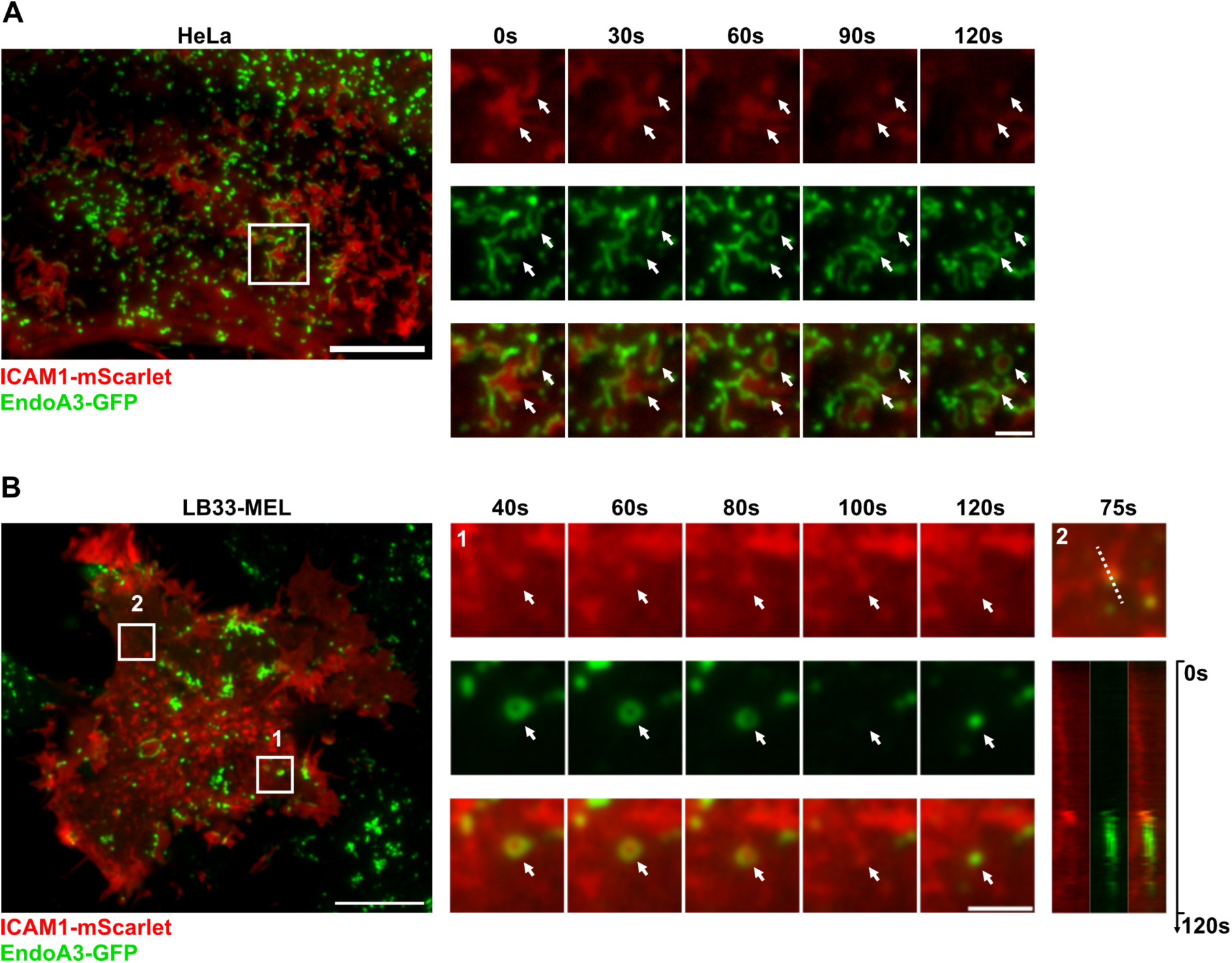
ICAM1 is an EndoA3-dependent endocytic cargo in cancer cells. A-B Supplementary representative live-cell TIRF images of EndoA3-GFP (stable) and ICAM1-mScarlet (transient) in HeLa (A) and LB33-MEL (B) cells. Time series show enlarged cropped areas extracted from Movie S5 (A) and Movie S6 (B). White arrowheads indicate dynamic co-distribution of both signals. Kymograph in (B) was made along dashed line in the enlarged cropped area corresponding to region 2 (Movie S7). Scale bars: 10 μm (full size image), 2 μm (enlarged cropped area). Data information: images are representative of two independent experiments.

**Fig. S3.**
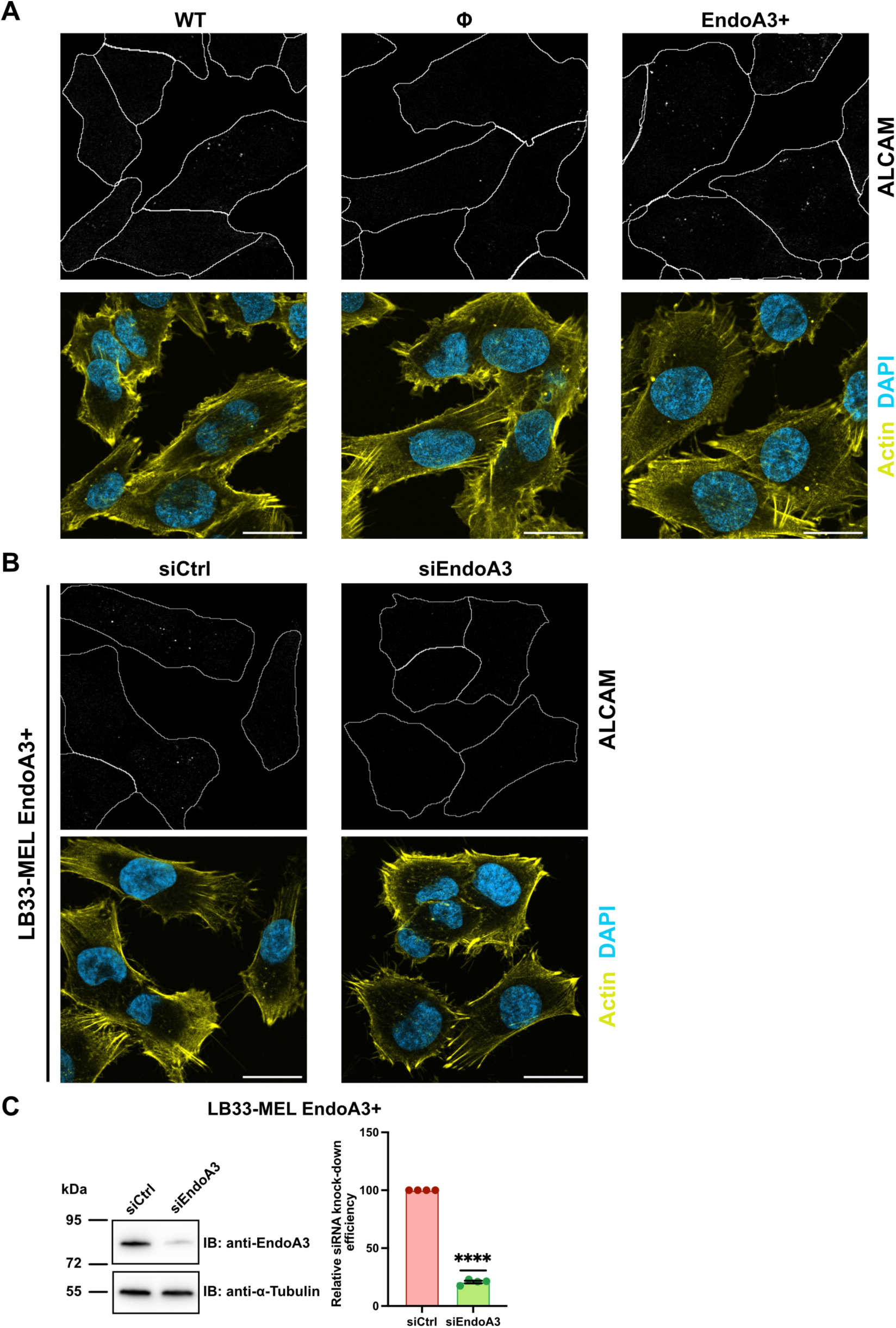
EndoA3-dependent CIE mediates the uptake of immune synapse components ALCAM in LB33-MEL cells. A-B Confocal images of anti-ALCAM (white spots) internalization in LB33-MEL cells stained for actin (phalloidin, yellow) and nuclei (DAPI, blue). In (A), wild-type (WT), stably transfected with empty plasmid (Φ), or stably expressing EndoA3-GFP (EndoA3+) LB33-MEL cells were used. In (B), LB33-MEL EndoA3+ cells were transfected with negative control (siCtrl) siRNA or EndoA3 targeting (siEndoA3) siRNA. Quantification for anti-ALCAM internalization shown in Fig. 2D-E. Scale bars: 20 μm. C Western blot analysis of LB33-MEL cells stably expressing EndoA3-GFP (EndoA3+ LB33-MEL) transfected for 72h with siRNAs: negative control (siCtrl) or against EndoA3 (siEndoA3). Immunodetection made with anti-EndoA3 and anti-α-Tubulin (loading control) antibodies. Quantification of immunoblots shows depletion efficiency of EndoA3 (histogram). Data information: In (A-B), images are representative of three independent experiments. In (C), Western blot images are representative of four independent experiments, from which quantitative data are pooled. Data are presented as mean ± SEM. ****P < 0.0001 (One sample *t* and Wilcoxon test). Source data are available online for this figure.

**Fig. S4.**
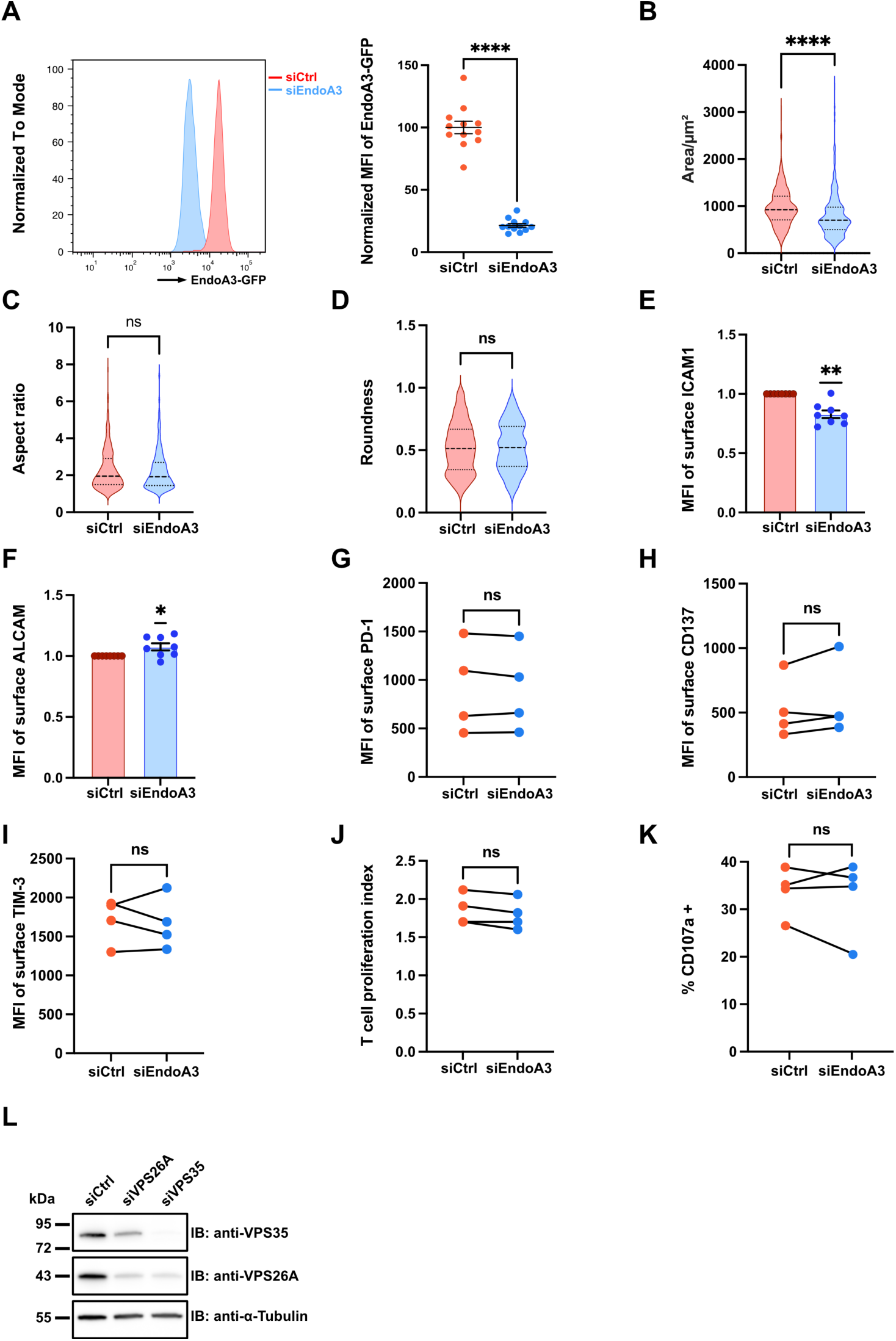
Effect of EndoA3 depletion on morphological parameters and surface levels of several markers in LB33-MEL EndoA3+ cells or CD8 T cells. A Flow cytometry analysis of LB33-MEL cells stably expressing EndoA3-GFP (LB33-MEL EndoA3+) transfected with siRNAs: negative control (siCtrl) or against EndoA3 (siEndoA3) (related to Fig. 3B-C). Quantification (scatter plot) based on GFP signal and presented as normalized MFI of GFP relative to siCtrl condition. B-D Quantification of physical parameters (surface area, aspect ratio and roundness) of LB33-MEL EndoA3+ cells transfected with siRNAs: negative control (siCtrl) or against EndoA3 (siEndoA3). E-F Quantification of surface ICAM1 (E) and ALCAM (F) levels on LB33-MEL EndoA3+ cells transfected with siRNAs: negative control (siCtrl) or against EndoA3 (siEndoA3). Quantified by flow cytometry analysis and presented as normalized MFI of ICAM1/ALCAM signal in siCtrl condition. G-I Quantification of T cell surface activation marker levels (PD-1, G; CD137, H; TIM-3, I) after 4 days of stimulation by LB33-MEL EndoA3+ cells transfected with either control siRNAs (siCtrl) or siRNAs against EndoA3 (siEndoA3). Quantified by flow cytometry analysis and presented as absolute MFI. J Quantification of CD8 T cell proliferation after 4 days of stimulation by LB33-MEL EndoA3+ cells transfected with either control siRNAs (siCtrl) or siRNAs against EndoA3 (siEndoA3). Quantified by flow cytometry analysis and presented as proliferation index. K Quantification of CD8 T cell degranulation percentage after a 10 min stimulation by LB33-MEL EndoA3+ cells transfected with either control siRNAs (siCtrl) or siRNAs against EndoA3 (siEndoA3). Quantified by flow cytometry analysis as the percentage of CD107a-positive cells. L Western blot analysis of LB33-MEL EndoA3+ cells transfected with siRNAs: negative control (siCtrl), or against the retromer subunits (siVPS35 or siVPS26A) (related to Fig. 3G). Immunodetection with anti-VPS35, anti-VPS26A and anti-α-Tubulin (loading control) antibodies. Data information: In (A), data are pooled from twelve independent experiments. In (B-D), data are pooled from three independent experiments. In (E-F), data are pooled from eight independent experiments. In (G-J), data are pooled from four independent experiments. In (B-D), data are presented as median and quartiles. In (A, E-F), data are presented as mean ± SEM. ns, not significant. *P < 0.05, **P < 0.01, ****P < 0.0001 (A and G-K, paired *t* test; B-D, Mann Whitney test; E-F, one sample *t* test). In (L), the Western blot image is representative of three independent experiments. Source data are available online for this figure.

**Fig. S5.**
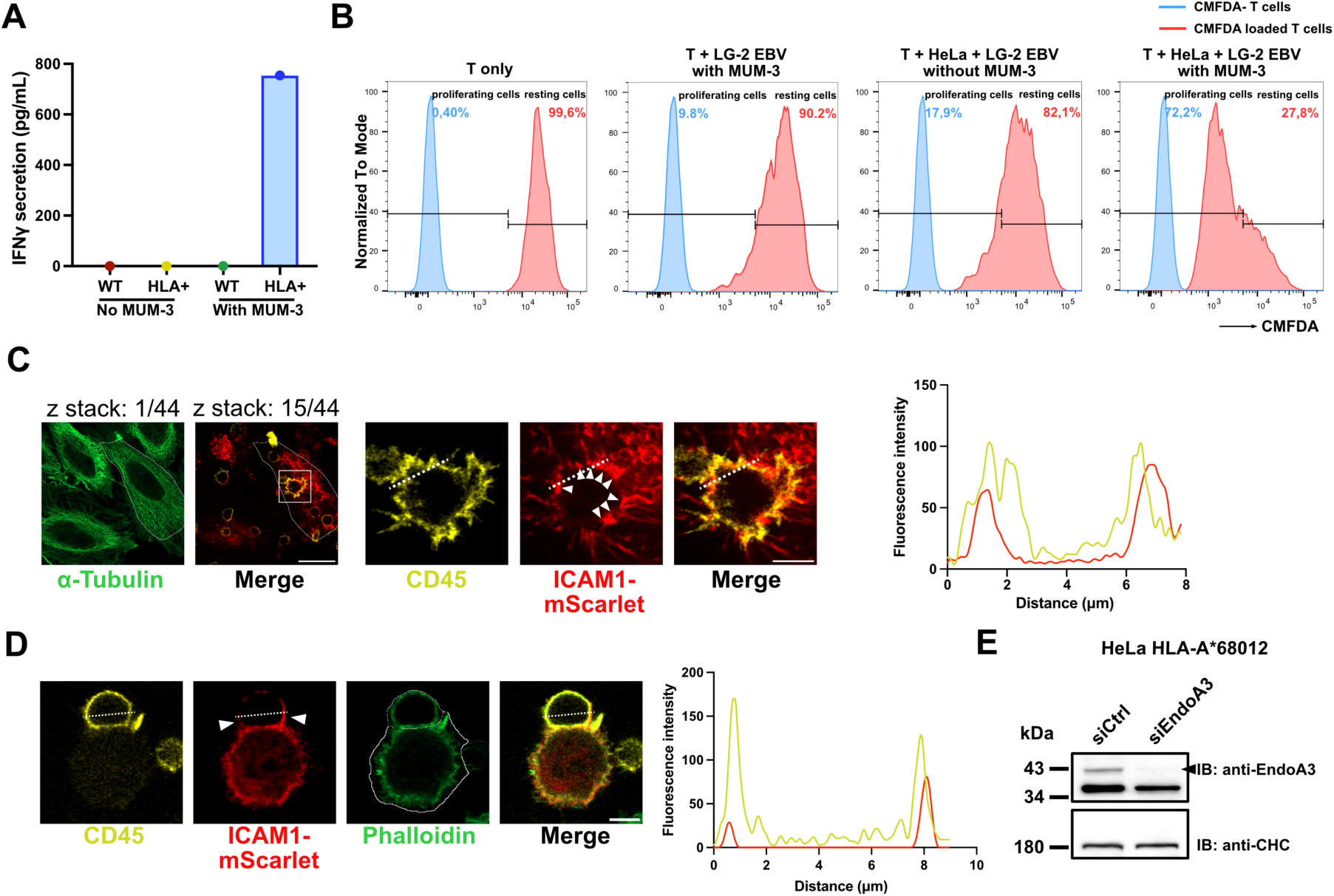
HLA-A*68012-expressing HeLa cells stimulate the activation of anti-MUM-3 CD8 T cells in the presence of MUM-3 peptide. A Quantification of IFNγ secretion (detected by ELISA) from anti-MUM-3 CD8 T cells, co-cultured for 20h with wild-type HeLa cells (WT) or stable HLA-A*68012-expressing HeLa cells (HLA+), with or without the presence of 300 ng/mL MUM-3 antigenic peptide. Data show absolute concentration of secreted IFNγ. B Flow cytometry analysis of anti-MUM-3 CD8 T cell proliferation after culture alone, or co-culture with LG-2 EBV feeder cells, or co-culture with both HLA-A*68012-expressing HeLa cells and LG-2 EBV feeder cells, with or without the presence of MUM-3 antigenic peptide, for 4 days. Proliferation was measured by the progressive dilution of CMFDA signal in T cells over generations. C Airyscan images of an immune synapse-like conjugate formed between a CD8 T cell (stained for CD45, yellow) and an adherent stable HLA-A*68012-expressing HeLa cell (stained for microtubules, green) transiently expressing ICAM1-mScarlet (red). White arrowheads indicate recruitment of ICAM1 to the vicinity of CD8 T cell. The fluorescence intensity profile was made along the dashed line region in enlarged cropped area. Scale bars: 20 μm (full size image), 5 μm (enlarged cropped areas). D Airyscan images of another immune synapse-like conjugate formed between a CD8 T cell (stained for CD45, yellow) and a stable HLA-A*68012-expressing HeLa cell (stained for actins, green) transiently expressing ICAM1-mScarlet (red) in suspension. The fluorescence intensity profile was made along the dashed line region. Cell segmentation (white contour) was made based on the actin staining for quantification (Fig. 5B). Scale bar: 5 μm. E Western blots analysis of stable HLA-A*68012-expressing HeLa cells transfected with siRNAs: negative control (siCtrl) or against EndoA3 (siEndoA3) (Fig. 5B). Immunodetection made with anti-EndoA3 and anti-clathrin heavy chain (CHC, loading control) antibodies. Data information: In (A-B), data and images are from single experiments. In (C-E), images are representative of three independent experiments.

**Fig. S6.**
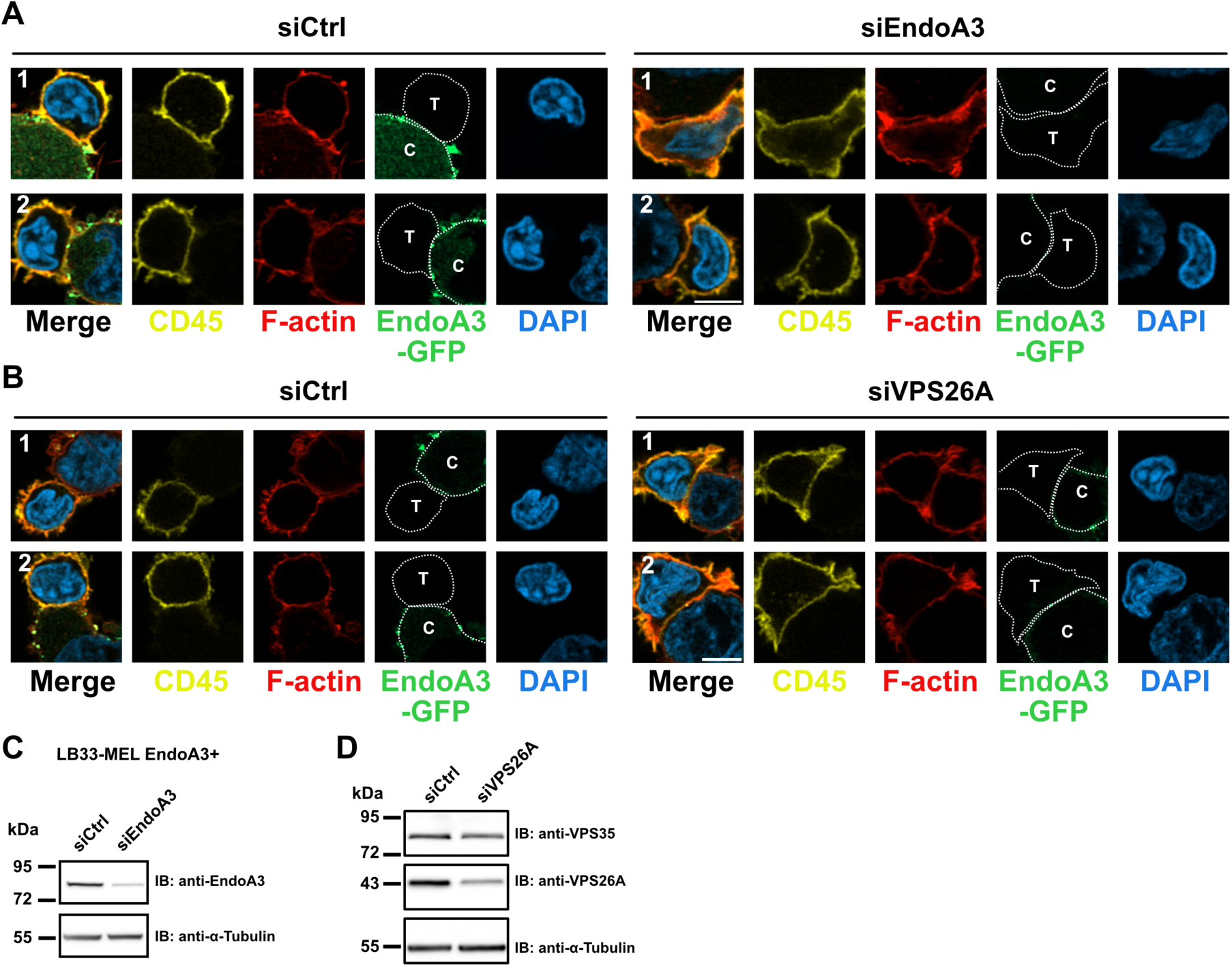
Inhibition of EndoA3-mediated endocytosis and retrograde transport causes enlarged immune synapses. A-B Separate channels of Airyscan images from Fig. 4D and 4F, respectively. CD45 (yellow), EndoA3 (green), actin (red) and nucleus (blue) are shown. C, cancer cell; T, CD8 T cell. Scale bar: 5 μm. C-D Western blot analyses of stable EndoA3-GFP-expressing LB33-MEL cells transfected with siRNAs: negative control (siCtrl), siRNAs against EndoA3 (siEndoA3) (Fig. 4E) or against VPS26A (siVPS26A) (Fig. 4G). In (C), immunodetection was made with an anti-EndoA3 antibody. In (D), immunodetection was made with anti-VPS35 and anti-VPS26A antibodies. In both panels, anti-α-Tubulin was used as a loading control. Data information: images are representative of three independent experiments.

**Movie S1 (separate file). Gradual disappearance of EndoA3-enclosed ICAM1 from the plasma membrane of a HeLa cell, related to Fig. 2A, cropped area 1.** HeLa cells stably expressing EndoA3-GFP (green) and transiently expressing ICAM1-mScarlet (red) were imaged for 2 minutes using live-cell TIRF microscopy, with 1-second intervals between frames. Representative of two independent experiments. Scale bar: 10 µm.

**Movie S2 (separate file). Gradual disappearance of EndoA3-enclosed ICAM1 from the plasma membrane of an LB33-MEL cell, related to Fig. 2B, cropped area 1.** LB33-MEL cells stably expressing EndoA3-GFP (green) and transiently expressing ICAM1-mScarlet (red) were imaged for 2 minutes using live-cell TIRF microscopy, with 1-second intervals between frames. Representative of two independent experiments. Scale bars: 10 μm (full size image), 2 μm (enlarged cropped area).

**Movie S3 (separate file). A punctate ICAM1-positive structure recruits EndoA3 and both signals disappear simultaneously from the plasma membrane of HeLa cell, related to Fig. 2A, kymograph of cropped area 2.** HeLa cells stably expressing EndoA3-GFP (green) and transiently expressing ICAM1-mScarlet (red) were imaged for 2 minutes using live-cell TIRF microscopy, with 1-second intervals between frames. The punctate ICAM1 structure is indicated by a white arrowhead. Representative of two independent experiments. Scale bar: 10 µm.

**Movie S4 (separate file). A punctate ICAM1-positive structure recruits EndoA3 and both signals disappear simultaneously from the plasma membrane of an LB33-MEL cell, related to Fig. 2B, kymograph of cropped area 2.** LB33-MEL cells stably expressing EndoA3-GFP (green) and transiently expressing ICAM1-mScarlet (red) were imaged for 2 minutes using live-cell TIRF microscopy, with 1-second intervals between frames. The punctate ICAM1 structure is indicated by a white arrowhead. Representative of two independent experiments. Scale bars: 10 μm (full size image), 2 μm (enlarged cropped area).

**Movie S5 (separate file). Dynamic ICAM1-positive patch, bordered by “lasso-like” EndoA3 structures, progressively shrinks and disappears from the plasma membrane of a HeLa cell, related to Fig. S2A.** HeLa cells stably expressing EndoA3-GFP (green) and transiently expressing ICAM1-mScarlet (red) were imaged for 2 minutes using live-cell TIRF microscopy, with 1-second intervals between frames. Representative of two independent experiments. Scale bars: 10 μm (full size image), 2 μm (enlarged cropped area).

**Movie S6 (separate file). Dynamic ICAM1-positive patch, bordered by “lasso-like” EndoA3 structures, progressively shrinks and disappears from the plasma membrane of an LB33-MEL cell, related to Fig. S2B, cropped area 1.** LB33-MEL cells stably expressing EndoA3-GFP (green) and transiently expressing ICAM1-mScarlet (red) were imaged for 2 minutes using live-cell TIRF microscopy, with 1-second intervals between frames. Representative of two independent experiments. Scale bar: 10 µm.

**Movie S7 (separate file). A punctate ICAM1-positive structure recruits EndoA3 and both signals disappear simultaneously from the plasma membrane of an LB33-MEL cell, related to Fig. S2B, kymograph of cropped area 2.** LB33-MEL cells stably expressing EndoA3-GFP (green) and transiently expressing ICAM1-mScarlet (red) were imaged for 2 minutes using live-cell TIRF microscopy, with 1-second intervals between frames. The punctate ICAM1 structure is indicated by a white arrowhead. Representative of two independent experiments. Scale bar: 10 µm.

**Movie S8 (separate file). ICAM1 tubulo-vesicular carrier flux is stronger and directed toward the contact region of an immune synapse-like conjugate formed between a HeLa cell and a T cell, related to Fig. 4**. A HeLa cell transiently expressing ICAM1-EGFP (green) forming an immune synapse-like conjugate with a CD8 T cell (red) was imaged by live-cell spinning-disk confocal microscopy, for 3 minutes, with 1-second intervals between frames. ICAM1-positive tubulo-vesicular carriers are observed fusing at the ICAM1-enriched contact zone, which grows and expands over time. Notably, anterograde transport of ICAM1-positive carriers toward the contact zone appears stronger than retrograde transport, and carrier density is higher in the region of the HeLa cell engaged in the immune synapse compared with regions not involved in synapse formation. Associated tracking and quantification data are shown in Fig. 4C-G. Representative of two independent experiments. Scale bar: 20 µm.

**Movie S9 (separate file). ICAM1 tubulo-vesicular carriers fuse at the ICAM1-enriched contact region of an immune synapse-like conjugate formed between a HeLa cell and a T cell, related to Fig. 4B (region 1’).** A HeLa cell transiently expressing ICAM1-EGFP (green) forming an immune synapse-like conjugate with a CD8 T cell (red) was imaged by live-cell spinning-disk confocal microscopy, for 3 minutes, with 1-second intervals between frames. ICAM1-positive tubulo-vesicular carriers are observed fusing at the ICAM1-enriched contact zone (white arrows), which grows and expands over time. Representative of two independent experiments. Scale bar: 5 µm.

## References

1. M. Kaksonen, A. Roux, Mechanisms of clathrin-mediated endocytosis. Nat Rev Mol Cell Biol 19, 313–326 (2018).

2. H. F. Renard, E. Boucrot, Unconventional endocytic mechanisms. Curr Opin Cell Biol 71, 120–129 (2021).

3. B. Sonnichsen, S. De Renzis, E. Nielsen, J. Rietdorf, M. Zerial, Distinct membrane domains on endosomes in the recycling pathway visualized by multicolor imaging of Rab4, Rab5, and Rab11. J Cell Biol 149, 901–914 (2000).

4. P. van der Sluijs et al., The small GTP-binding protein rab4 controls an early sorting event on the endocytic pathway. Cell 70, 729–740 (1992).

5. M. N. Seaman, Cargo-selective endosomal sorting for retrieval to the Golgi requires retromer. J Cell Biol 165, 111–122 (2004).

6. K. E. McNally et al., Retriever is a multiprotein complex for retromer-independent endosomal cargo recycling. Nat Cell Biol 19, 1214–1225 (2017).

7. M. D. Healy et al., Structure of the endosomal Commander complex linked to Ritscher-Schinzel syndrome. Cell 186, 2219–2237 e2229 (2023).

8. M. N. Seaman, J. M. McCaffery, S. D. Emr, A membrane coat complex essential for endosome-to-Golgi retrograde transport in yeast. J Cell Biol 142, 665–681 (1998).

9. M. Shafaq-Zadah, E. Dransart, L. Johannes, Clathrin-independent endocytosis, retrograde trafficking, and cell polarity. Curr Opin Cell Biol 65, 112–121 (2020).

10. M. Shafaq-Zadah et al., Persistent cell migration and adhesion rely on retrograde transport of beta(1) integrin. Nat Cell Biol 18, 54–64 (2016).

11. J. M. Carpier et al., Rab6-dependent retrograde traffic of LAT controls immune synapse formation and T cell activation. J Exp Med 215, 1245–1265 (2018).

12. N. M. Dieckmann, G. L. Frazer, Y. Asano, J. C. Stinchcombe, G. M. Griffiths, The cytotoxic T lymphocyte immune synapse at a glance. J Cell Sci 129, 2881–2886 (2016).

13. B. J. Peter et al., BAR Domains as Sensors of Membrane Curvature: The Amphiphysin BAR Structure. SCIENCE 303 (2004).

14. J. L. Gallop et al., Mechanism of endophilin N-BAR domain-mediated membrane curvature. EMBO J 25, 2898–2910 (2006).

15. A. B. Sparks, N. G. Hoffman, S. J. McConnell, D. M. Fowlkes, B. K. Kay, Cloning of ligand targets: systematic isolation of SH3 domain-containing proteins. Nat Biotechnol 14, 741–744 (1996).

16. K. D. Micheva, B. K. Kay, P. S. McPherson, Synaptojanin forms two separate complexes in the nerve terminal. Interactions with endophilin and amphiphysin. J Biol Chem 272, 27239–27245 (1997).

17. C. Giachino et al., A novel SH3-containing human gene family preferentially expressed in the central nervous system. Genomics 41, 427–434 (1997).

18. C. W. So et al., Expression and protein-binding studies of the EEN gene family, new interacting partners for dynamin, synaptojanin and huntingtin proteins. Biochem J 348 **Pt 2**, 447–458 (2000).

19. N. Ringstad, Y. Nemoto, P. De Camilli, The SH3p4/Sh3p8/SH3p13 protein family: binding partners for synaptojanin and dynamin via a Grb2-like Src homology 3 domain. Proc Natl Acad Sci U S A 94, 8569–8574 (1997).

20. F. Simpson et al., SH3-domain-containing proteins function at distinct steps in clathrin-coated vesicle formation. Nat Cell Biol 1, 119–124 (1999).

21. H. Gad et al., Fission and Uncoating of Synaptic Clathrin-Coated Vesicles Are Perturbed by Disruption of Interactions with the SH3 Domain of Endophilin. Neuron 27, 301–312 (2000).

22. I. Milosevic et al., Recruitment of endophilin to clathrin-coated pit necks is required for efficient vesicle uncoating after fission. Neuron 72, 587–601 (2011).

23. E. Boucrot et al., Endophilin marks and controls a clathrin-independent endocytic pathway. Nature 517, 460–465 (2015).

24. H. F. Renard et al., Endophilin-A2 functions in membrane scission in clathrin-independent endocytosis. Nature 517, 493–496 (2015).

25. M. Simunovic et al., Friction Mediates Scission of Tubular Membranes Scaffolded by BAR Proteins. Cell 170, 172–184 e111 (2017).

26. H. F. Renard et al., Endophilin-A3 and Galectin-8 control the clathrin-independent endocytosis of CD166. Nat Commun 11, 1457 (2020).

27. L. Johannes, C. Wunder, M. Shafaq-Zadah, Glycolipids and Lectins in Endocytic Uptake Processes. J Mol Biol 428, 4792–4818 (2016).

28. R. Lakshminarayan et al., Galectin-3 drives glycosphingolipid-dependent biogenesis of clathrin-independent carriers. Nat Cell Biol 16, 595–606 (2014).

29. C. Lemaigre et al., N-BAR and F-BAR proteins-endophilin-A3 and PSTPIP1-control clathrin-independent endocytosis of L1CAM. Traffic 24, 190–212 (2023).

30. F. Arai, O. Ohneda, T. Miyamoto, X. Q. Zhang, T. Suda, Mesenchymal stem cells in perichondrium express activated leukocyte cell adhesion molecule and participate in bone marrow formation. J Exp Med 195, 1549–1563 (2002).

31. F. Ferragut, V. S. Vachetta, M. F. Troncoso, G. A. Rabinovich, M. T. Elola, ALCAM/CD166: A pleiotropic mediator of cell adhesion, stemness and cancer progression. Cytokine Growth Factor Rev 61, 27–37 (2021).

32. W. Weichert, T. Knosel, J. Bellach, M. Dietel, G. Kristiansen, ALCAM/CD166 is overexpressed in colorectal carcinoma and correlates with shortened patient survival. J Clin Pathol 57, 1160–1164 (2004).

33. L. C. van Kempen et al., Molecular basis for the homophilic activated leukocyte cell adhesion molecule (ALCAM)-ALCAM interaction. J Biol Chem 276, 25783–25790 (2001).

34. M. A. Bowen et al., Cloning, mapping, and characterization of activated leukocyte-cell adhesion molecule (ALCAM), a CD6 ligand. J Exp Med 181, 2213–2220 (1995).

35. M. Kamoun, M. E. Kadin, P. J. Martin, J. Nettleton, J. A. Hansen, A novel human T cell antigen preferentially expressed on mature T cells and shared by both well and poorly differentiated B cell leukemias and lymphomas. J Immunol 127, 987–991 (1981).

36. A. Aruffo et al., CD6-ligand interactions: a paradigm for SRCR domain function? Immunol Today 18, 498–504 (1997).

37. A. W. Zimmerman et al., Long-term engagement of CD6 and ALCAM is essential for T-cell proliferation induced by dendritic cells. Blood 107, 3212–3220 (2006).

38. N. J. Hassan, A. N. Barclay, M. H. Brown, Frontline: Optimal T cell activation requires the engagement of CD6 and CD166. Eur J Immunol 34, 930–940 (2004).

39. I. Gimferrer et al., Relevance of CD6-mediated interactions in T cell activation and proliferation. J Immunol 173, 2262–2270 (2004).

40. G. A. Van Seventer, Y. Shimizu, K. J. Horgan, S. Shaw, The LFA-1 ligand ICAM-1 provides an important costimulatory signal for T cell receptor-mediated activation of resting T cells. J Immunol 144, 4579–4586 (1990).

41. L. A. Zuckerman, L. Pullen, J. Miller, Functional consequences of costimulation by ICAM-1 on IL-2 gene expression and T cell activation. J Immunol 160, 3259–3268 (1998).

42. M. J. Deeths, M. F. Mescher, ICAM-1 and B7-1 provide similar but distinct costimulation for CD8+ T cells, while CD4+ T cells are poorly costimulated by ICAM-1. European Journal of Immunology 29, 45–53 (1999).

43. L. Johannes, M. Shafaq-Zadah, SNAP-tagging the retrograde route. Methods Cell Biol 118, 139–155 (2013).

44. M. Shafaq-Zadah, E. Dransart, L. Johannes, Quantitative Methods to Study Endocytosis and Retrograde Transport of Cargo Proteins. Methods Mol Biol 2233, 53–70 (2021).

45. F. Lehmann et al., Differences in the antigens recognized by cytolytic T cells on two successive metastases of a melanoma patient are consistent with immune selection. Eur J Immunol 25, 340–347 (1995).

46. A. Hierro et al., Functional architecture of the retromer cargo-recognition complex. Nature 449, 1063–1067 (2007).

47. A. Bugarcic et al., Vps26A and Vps26B subunits define distinct retromer complexes. Traffic 12, 1759–1773 (2011).

48. M. C. Kerr et al., A novel mammalian retromer component, Vps26B. Traffic 6, 991–1001 (2005).

49. L. Fourriere et al., RAB6 and microtubules restrict protein secretion to focal adhesions. J Cell Biol 218, 2215–2231 (2019).

50. F. Mallard et al., Early/recycling endosomes-to-TGN transport involves two SNARE complexes and a Rab6 isoform. J Cell Biol 156, 653–664 (2002).

51. F. Tyckaert, N. Zanin, P. Morsomme, H. F. Renard, Rac1, the actin cytoskeleton and microtubules are key players in clathrin-independent endophilin-A3-mediated endocytosis. J Cell Sci 135 (2022).

52. J. F. Baurain et al., High frequency of autologous anti-melanoma CTL directed against an antigen generated by a point mutation in a new helicase gene. J Immunol 164, 6057–6066 (2000).

53. O. Kjaerulff, L. Brodin, A. Jung, The structure and function of endophilin proteins. Cell Biochem Biophys 60, 137–154 (2011).

54. L. M. Traub, J. S. Bonifacino, Cargo recognition in clathrin-mediated endocytosis. Cold Spring Harb Perspect Biol 5, a016790 (2013).

55. P. Moreno-Layseca et al., Cargo-specific recruitment in clathrin- and dynamin-independent endocytosis. Nat Cell Biol 23, 1073–1084 (2021).

56. Y. Tang et al., Identification of the endophilins (SH3p4/p8/p13) as novel binding partners for the beta1-adrenergic receptor. Proc Natl Acad Sci U S A 96, 12559–12564 (1999).

57. P. Soubeyran, K. Kowanetz, I. Szymkiewicz, W. Y. Langdon, I. Dikic, Cbl-CIN85-endophilin complex mediates ligand-induced downregulation of EGF receptors. Nature 416, 183–187 (2002).

58. A. Petrelli et al., The endophilin-CIN85-Cbl complex mediates ligand-dependent downregulation of c-Met. Nature 416, 187–190 (2002).

59. G. Genet et al., Endophilin-A2 dependent VEGFR2 endocytosis promotes sprouting angiogenesis. Nat Commun 10, 2350 (2019).

60. F. Steinberg et al., A global analysis of SNX27-retromer assembly and cargo specificity reveals a function in glucose and metal ion transport. Nat Cell Biol 15, 461–471 (2013).

61. J. H. Jo et al., Recycling and LFA-1-dependent trafficking of ICAM-1 to the immunological synapse. J Cell Biochem 111, 1125–1137 (2010).

62. A. L. Mallam, E. M. Marcotte, Systems-wide Studies Uncover Commander, a Multiprotein Complex Essential to Human Development. Cell Syst 4, 483–494 (2017).

63. C. Wan et al., Panorama of ancient metazoan macromolecular complexes. Nature 525, 339–344 (2015).

64. A. Singla et al., Endosomal PI(3)P regulation by the COMMD/CCDC22/CCDC93 (CCC) complex controls membrane protein recycling. Nat Commun 10, 4271 (2019).

65. K. E. Chen, M. D. Healy, B. M. Collins, Towards a molecular understanding of endosomal trafficking by Retromer and Retriever. Traffic 20, 465–478 (2019).

66. A. Fuse et al., VPS29-VPS35 intermediate of retromer is stable and may be involved in the retromer complex assembly process. FEBS Lett 589, 1430–1436 (2015).

67. M. A. Purbhoo, D. J. Irvine, J. B. Huppa, M. M. Davis, T cell killing does not require the formation of a stable mature immunological synapse. Nat Immunol 5, 524–530 (2004).

68. M. R. Jenkins et al., Failed CTL/NK cell killing and cytokine hypersecretion are directly linked through prolonged synapse time. J Exp Med 212, 307–317 (2015).

69. A. J. Davenport et al., CAR-T Cells Inflict Sequential Killing of Multiple Tumor Target Cells. Cancer Immunol Res 3, 483–494 (2015).

70. A. J. Davenport et al., Chimeric antigen receptor T cells form nonclassical and potent immune synapses driving rapid cytotoxicity. Proc Natl Acad Sci U S A 115, E2068–E2076 (2018).

71. M. Cazaux et al., Single-cell imaging of CAR T cell activity in vivo reveals extensive functional and anatomical heterogeneity. J Exp Med 216, 1038–1049 (2019).

72. A. E. Petit et al., A major secretory defect of tumour-infiltrating T lymphocytes due to galectin impairing LFA-1-mediated synapse completion. Nat Commun 7, 12242 (2016).

73. G. Shi et al., SNAP-tag based proteomics approach for the study of the retrograde route. Traffic 13, 914–925 (2012).

74. G. Degiovanni, T. Lahaye, M. Hérin, P. Hainaut, T. Boon, Antigenic heterogeneity of a human melanoma tumor detected by autologous CTL clones. European Journal of Immunology 18, 671–676 (1988).

75. C. Stringer, M. Pachitariu, Cellpose3: one-click image restoration for improved cellular segmentation. bioRxiv 10.1101/2024.02.10.579780 (2024).

76. I. Arganda-Carreras et al., Trainable Weka Segmentation: a machine learning tool for microscopy pixel classification. Bioinformatics 33, 2424–2426 (2017).

77. J. Y. Tinevez et al., TrackMate: An open and extensible platform for single-particle tracking. Methods 115, 80–90 (2017).

78. C. Stringer, T. Wang, M. Michaelos, M. Pachitariu, Cellpose: a generalist algorithm for cellular segmentation. Nat Methods 18, 100–106 (2021).

